# Differential Phagocytosis induces Diverse Macrophage Activation States in Malignant Gliomas

**DOI:** 10.1101/2025.03.15.642920

**Authors:** Senthilnath Lakshmanachetty, Kent Riemondy, Bridget Sanford, Andrew Donson, Ilango Balakrishnan, Eric Prince, Todd Hankinson, Nathan Dahl, Rajeev Vibhakar, Nicholas K. Foreman, Sujatha Venkataraman, Siddhartha S. Mitra

## Abstract

Diffuse midline glioma (DMG) and Glioblastoma are malignant brain tumors in pediatric and adult patients. The current standard-of-care treatment for DMG is radiotherapy (RT), whereas GBM treatment includes surgery, followed by RT and chemotherapy. Although RT is known to modulate immune responses in cancer and enhance the effectiveness of myeloid checkpoint blockade, the downstream macrophage responses to differential phagocytosis induction remain poorly understood. This study examined macrophage-mediated phagocytosis caused by either RT, anti-CD47 checkpoint blockade, or their combination. We found that RT increased the expression of several damage-associated molecular patterns on the surface of glioma cell lines. Furthermore, RT enhanced anti-CD47-mediated macrophage phagocytosis of glioma cell lines *in vitro*. Single-cell RNA-sequencing revealed the diverse transcriptional and functional signatures of human macrophage subsets that either promoted or inhibited phagocytosis of glioma cells pretreated with RT, anti-CD47 therapy, or both. Consistent with these results, the combination therapy significantly reduced tumor growth, prolonged survival in glioma-bearing mice, and induced distinct macrophage activation states *in vivo* compared to either treatment alone. These findings highlight the plasticity and heterogeneity of macrophage responses during phagocytosis and provide compelling evidence for combining RT with anti-CD47 therapy as a promising therapeutic strategy for glioma treatment.

## Introduction

Diffuse intrinsic pontine glioma (DIPG) represents a category of diffuse midline gliomas that originate in the pons area during childhood. The outlook for DIPG patients is extremely poor, as over 90% succumb to the disease within two years following diagnosis, with the median survival period ranging from 9 to 12 months^1^. The strategic location in the brainstem and the tumor’s tendency to spread into nearby healthy tissue greatly restrict the options for surgical intervention, positioning DIPG as one of the most dire and complex cancers affecting children. Currently, radiotherapy (RT) is the primary treatment for DIPG, though it does not cure the disease. However, it offers temporary relief from symptoms and a slight extension in life expectancy, adding about 3 months to the survival rate^2^. This fact emphasizes the critical need for new therapeutic approaches for DIPG. Although immune checkpoint inhibitors (ICIs) have been successful in treating various solid tumors, they have not been effective in pediatric high-grade glioma (pHGG) treatment^3^. This lack of efficacy is likely due to the unique environment of the DIPG tumor, which is characterized by a low presence of T cells and a predominance of microglia/macrophages^4, 5^. On the other hand, the immune microenvironment of adult glioblastoma (GBM) comprises macrophages/microglia, dendritic cells, myeloid-derived suppressor cells, and T cells^6^.

Phagocytosis is a critical function of the immune system, involving immune cells that encapsulate and break down large external pathogens or dying cells to preserve the body’s balance^7^. Macrophages play a pivotal role in this process by detecting signals on cells undergoing apoptosis or cancerous cells that indicate they should be engulfed and digested^8^. These signals include “find-me” and “eat-me” cues that trigger the phagocytic process^9^. Conversely, healthy cells and some cancer cells can emit “don’t-eat-me” signals that deter phagocytic activity^10^. A key interaction in this mechanism is between CD47 on the cell surface and SIRPα on macrophages, leading to the activation of processes that facilitate phagocytosis, such as cytoskeletal changes and membrane movement^11^. Beyond the CD47-SIRPα interaction^12^, an increasing number of signals have been identified that inhibit phagocytosis, including interactions between CD24-Siglec 10^13^, PD-L1-PD-1^14^, MHC-1-LILRB1^15^, and APMAP-GPR84^16^. Given that tumors often contain high levels of macrophages, inhibiting these “don’t-eat-me” signals presents a promising approach for enhancing cancer immunotherapy. Despite the progress in developing targeted therapies for various cancers, RT continues to be a cornerstone in cancer management, with over half of all cancer patients undergoing RT at some point during their illness. RT primarily works by causing immediate damage to the DNA of cancer cells, leading to their death^17^. Innovations such as intensity-modulated RT and image-guided RT have improved the precision of treatment, allowing for increased radiation doses to be delivered to tumors while minimizing exposure to adjacent healthy tissues^18^. Consequently, RT has become safer and more effective for a broader range of patients. Additionally, the destruction of cancer cells by RT releases tumor antigens, potentially enhancing the activation of antigen-specific T cells and triggering an adaptive immune response to target residual cancer cells^19^. This potential has sparked numerous clinical trials exploring the synergy between radiotherapy and immunotherapy^20^. However, radiotherapy’s impact on the immune system is complex, as it can provoke both protective and harmful immune responses, influencing tumor growth in varying ways.

In this study, we examined the effects of combining radiation therapy (RT) and CD47 blockade in pediatric diffuse midline glioma (DMG) and adult glioblastoma (GBM). Our findings reveal that RT induces immunogenic cell death in gliomas and amplifies macrophage-mediated phagocytosis of glioma cells when combined with CD47 blockade. Using single-cell RNA sequencing, we identified distinct macrophage subpopulations that either promoted or inhibited the phagocytosis of glioma cells pretreated with RT, anti-CD47 therapy, or both. Furthermore, the combination therapy significantly improved survival in several glioma mouse models by enhancing diverse macrophage activation states and tumor-killing activity. Collectively, our study sheds light on the heterogeneity and plasticity of macrophages during phagocytosis induced by RT or CD47 blockade and supports the use of this combination therapy in treating malignant gliomas.

## Results

### Radiotherapy induces immunogenic cell death in malignant gliomas

Radiotherapy (RT) is known to cause immunogenic cell death (ICD) in different cancer types^21–23^ including adult GBM^24, 25^. However, it is unclear if RT can elicit ICD in pediatric malignant gliomas (H3K27M-DMG and GBM). To address this question, we exposed several patient-derived human DMG (BT-245, SU-DIPGXVII, SU-DIPGXXV) and mouse glioma cell lines (SB28 and CT-2A) to increasing doses of RT for three consecutive days (0 Gy, 2 Gy X 3, 4Gy X 3, 8Gy X 3, 16 Gy X 3) and evaluated their response. We find the IC_50_ of human DMG lines to range from 10 to 12 Gy (BT-245, SU-DIPGXVII, SU-DIPGXXV), whereas the IC_50_ for mouse glioma lines were 9.3 Gy (CT-2A) and 14.2 Gy (SB28) (**Figure S1**). The cells undergoing ICD are characterized by the increased expression of damage-associated molecular patterns (DAMPs)^26^. We, therefore measured the surface expression of common DAMP molecules such as Phosphatidylserine (PS), Calreticulin (CRT), heat shock protein 70 (HSP70) and 90 (HSP90)^27^ at 24 hours after the last treatment with RT. In both the human DMG and mouse glioma cell lines, there is a dose-dependent increase in the expression of PS (**Figure 1A and Figure S2A**), CRT (**Figure 1B and Figure S2B**), HSP70 (**Figure 1C and Figure S2C**), and HSP90 (**Figure 1D and Figure S2D**) compared to non-irradiated (0 Gy) cells. We further measured the extracellular high mobility group protein 1 (HMGB1), a DAMP molecule released by dying cells^28^ in the supernatants of DMG and GBM cell lines that were collected at 48 hours after the last treatment with RT. In line with the findings for the surface expression of DAMPs, we observed a dose-dependent increase in the release of HMGB1 (**Figure 1E and Figure S2E**) compared to non-irradiated cell lines. Collectively, these findings indicate that fractionated RT induces ICD in both DMG and GBM cell lines as demonstrated by a dose-dependent increase in the surface expression and extracellular release of DAMPs.

**Figure 1:**
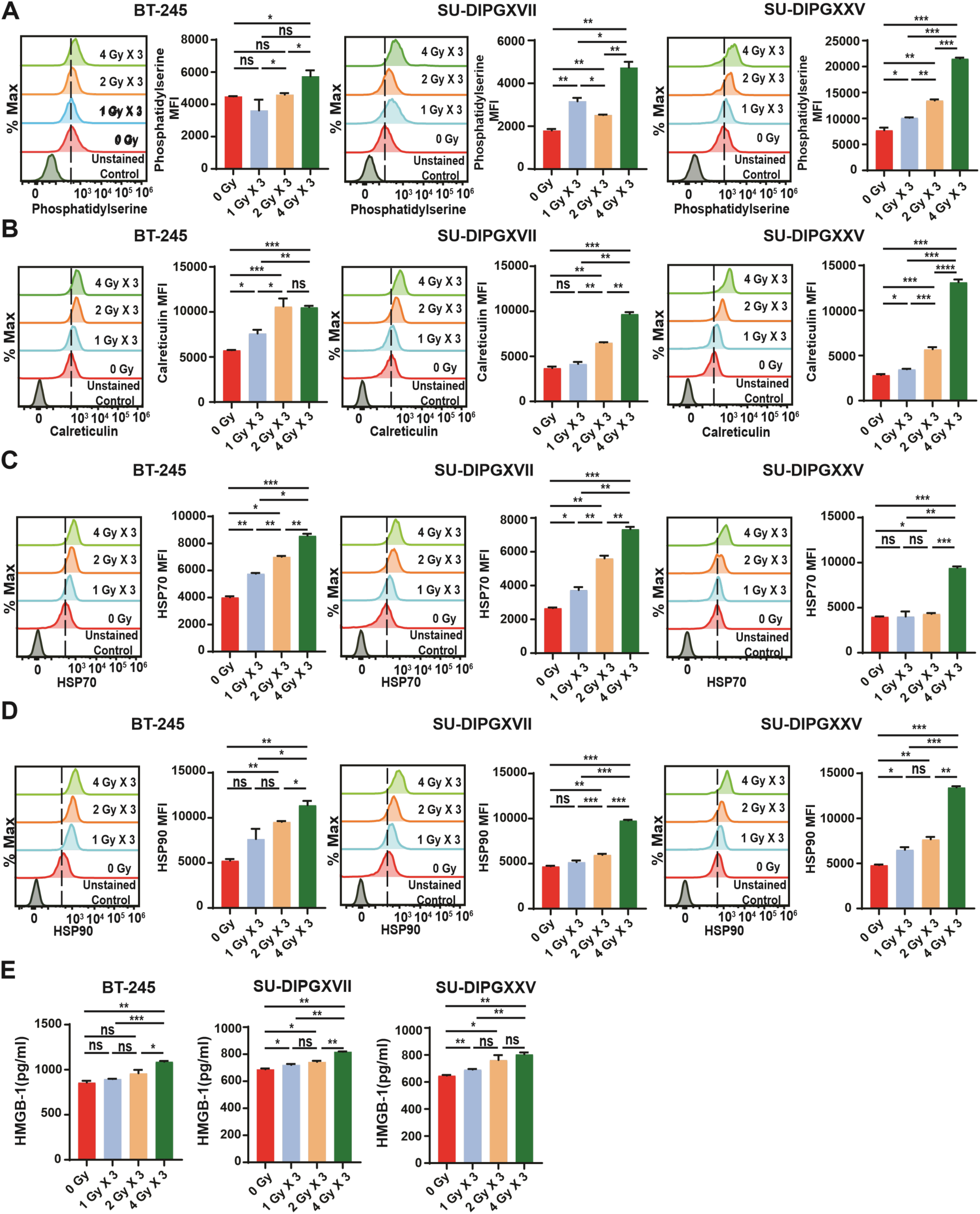
Radiation therapy induces the surface expression and release of damage-associated molecular patterns (DAMPs) in human DMG/DIPG. Human patient-derived DMG/DIPG cell lines were exposed to increasing doses of RT for 3 consecutive days. **(A-D)** Representative overlay histograms and median fluorescence intensity (MFI) values displaying the expression levels of phosphatidylserine (PS), Calreticulin (CRT), Heat shock protein (HSP70), and HSP 90 on the surface of BT245, SU-DIPGXVII and SU-DIPGXXV cells. Results are expressed as mean ± SD (n=3 technical replicates). ns, not significant. Unpaired student’s t-test: p < 0.05, **p < 0.01. **(E)** 24 hours post the final RT treatment, HMGB1 released in the supernatants of BT245, SU-DIPGXVII and SU-DIPGXXV cells were quantified. Results are expressed as mean ± SD (n=3 technical replicates). ns, not significant; Unpaired student’s t-test: p < 0.05, **p < 0.01.

### A combination of radiotherapy and anti-CD47 therapy enhances the *in vitro* phagocytosis of malignant glioma cell lines

DAMPs are known to activate the cells of the innate immune system and enhance adaptive immune responses either directly or indirectly^29, 30^. To examine if the increased expression of DAMPs on DMG and GBM cell lines can promote the clearance of DMG and GBM cell lines by macrophages, we performed *in vitro* phagocytosis assay. Briefly, the DMG cell lines were exposed to fractionated RT as described before and co-cultured with human peripheral blood mononuclear cell (PBMC)-derived macrophages at a 2:1 ratio in the presence or absence of blocking human anti-CD47 mAb. The phagocytosis of tumor cells by macrophages were determined by flow cytometry as previously published^31^ (**Figure S3)**. Anti-CD47 mAb treatment increased the phagocytosis of three different DMG cell lines compared to IgG treatment (**Figure 2A-2F**). However, RT (4 Gy X 3) significantly increased the phagocytosis of SU-DIPG25, but not SU-DIPG17 and BT-245 compared to control (**Figure 2A-2F**). Combining RT (4 Gy X 3) and anti-CD47 treatment significantly enhanced the phagocytosis of DMG cells compared to individual treatments (**Figure 2A-2F**). We further assessed the phagocytosis of mouse glioma cells that were either irradiated or non-irradiated and co-cultured with mouse bone marrow-derived macrophages in the presence or absence of anti-msCD47 mAb (clone MIAP301). Anti-CD47 treatment significantly increased the phagocytosis of SB28 cells, but not CT-2A cells compared to control (**Figure 2H-2J**). RT did not increase the phagocytosis of either SB28 or CT-2A cells (**Figure 2H-2J**). However, combining RT (3 Gy X 3) and anti-CD47 mAb treatment significantly enhanced the phagocytosis of both CT-2A and SB28 compared to either control, anti-CD47 mAb or RT treatment alone (**Figure 2H-2J**). These findings suggest that a combination of fractionated RT and anti-CD47 treatment significantly promotes the in vitro phagocytosis of human DMG/DIPG and mouse glioma cell lines. Notably, this combination enabled consistent phagocytosis even in cell lines that invariably escaped monotherapy.

**Figure 2:**
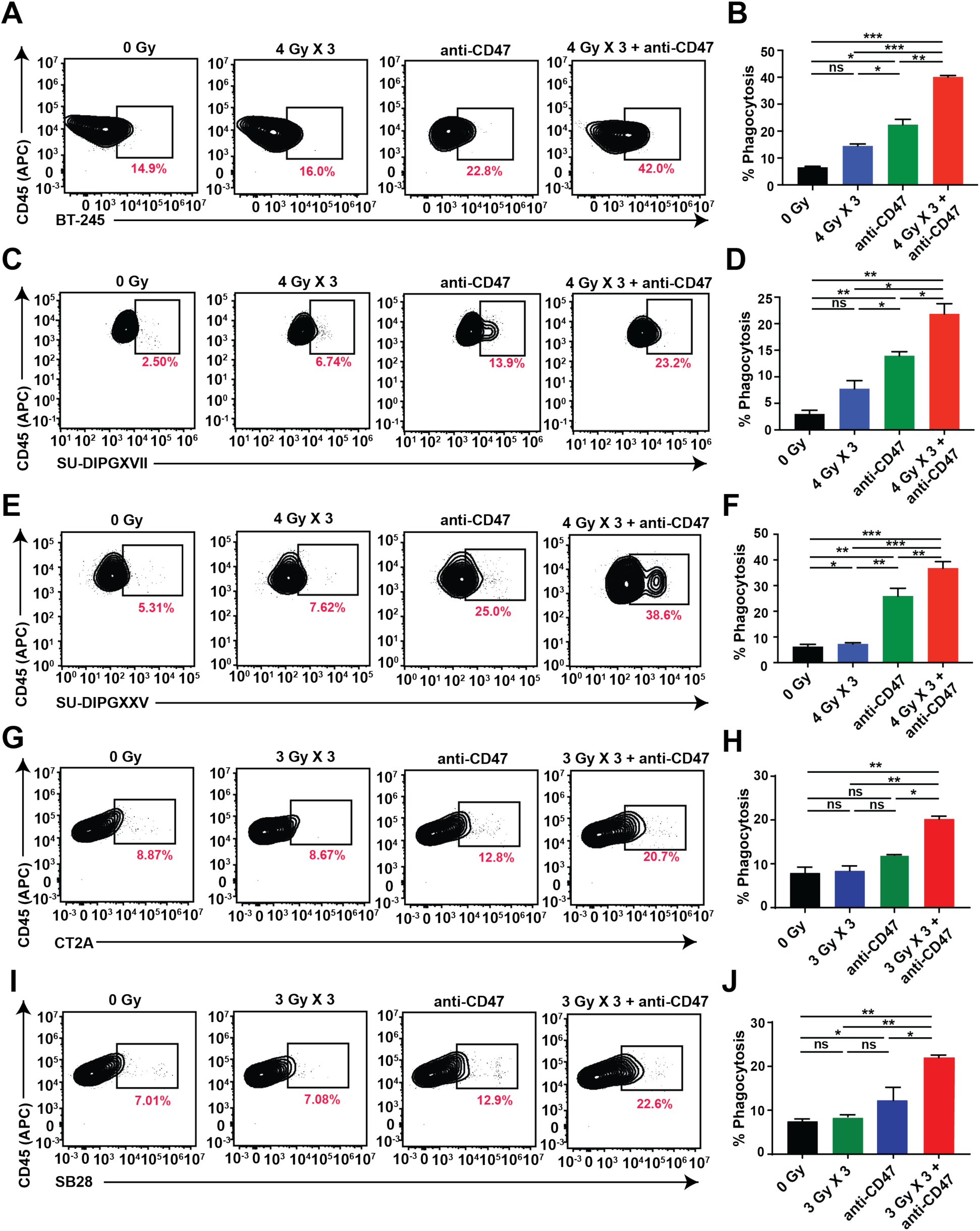
Combining radiation therapy with anti-CD47 mAb treatment enhances *in vitro* phagocytosis of human DIPG and mouse glioma cell lines. Human patient-derived DIPG cell lines (BT-245, SU-DIPGXVII, and SU-DIPGXXV) were exposed to either 0 or 4 Gy X 3 and incubated with human peripheral blood-derived macrophages in the presence of anti-CD47 mAb, HU5F9-G4. Flow cytometry (A, C, and E) as well as histogram (B, D, and F) plots show that combining fractionated irradiation and anti-CD47 antibody treatment increases the phagocytosis of DIPG/DMG cells by macrophages compared to individual treatments alone. **(G-J)** Mouse GBM cell lines (SB28 and CT2A) were exposed to either 0 or 4Gy X 3 and incubated with mouse bone marrow-derived macrophages in the presence of anti-CD47 mAb, MIAP301. Data shown are consistent with two independent experiments (n=3) and are shown as mean ± SD. ns, not significant; Unpaired student’s t-test. *p < 0.05, **p < 0.01 and ***p < 0.0001. One-way ANOVA, ****p < 0.0001 (BT245), ***p<0.001 (DIPGXVII) and ****p < 0.0001 (DIPGXXV). ***p < 0.001 (CT2A) and **p < 0.001 (SB28).

### Single-Cell RNA-Seq Identifies Distinct Macrophage Subsets Induced by DMG Cells Following Radiotherapy and Anti-CD47 Treatment

To understand the response of macrophages to different phagocytic stimuli, we performed an unbiased and comprehensive overview of the macrophages that had phagocytosed (“eaters”) tumor cells vs those that had not (“non-eaters”) in response to either control, RT, anti-CD47, or combo treatments. To this end, we co-cultured PBMC-derived macrophages with either irradiated (4 Gy X 3) or non-irradiated (O Gy) DMG (BT-245) cells in the presence or absence of anti-CD47 mAb for 24 hours and physically segregated the “eaters” from the “non-eaters” using flow cytometry and subjected them to single-cell RNA-sequencing (scRNA-Seq) as shown in **Figure 3A**. As expected from our earlier results, we observed more “eaters” and less “non-eaters” in RT plus anti-CD47 treatment (**Figure S4D**) compared to either control or monotherapy treatments (**Figure S4A-S4C**). In total, for the “eaters,” we obtained 6641 cells from the control, 2790 cells from the RT, 2014 cells from the anti-CD47, and 5732 cells from the RT plus anti-CD47 (**Figure 3D**) after performing quality control and filtering-out low-quality cells from scRNA-Seq datasets (see Materials and Methods). Dimension reduction and clustering of the cells based on their gene expression profiles identified 11 different clusters (C0-C11) (**Figure 3B**). Of these, eight were macrophage clusters and three were identified as contaminating tumor cell clusters.

**Figure 3:**
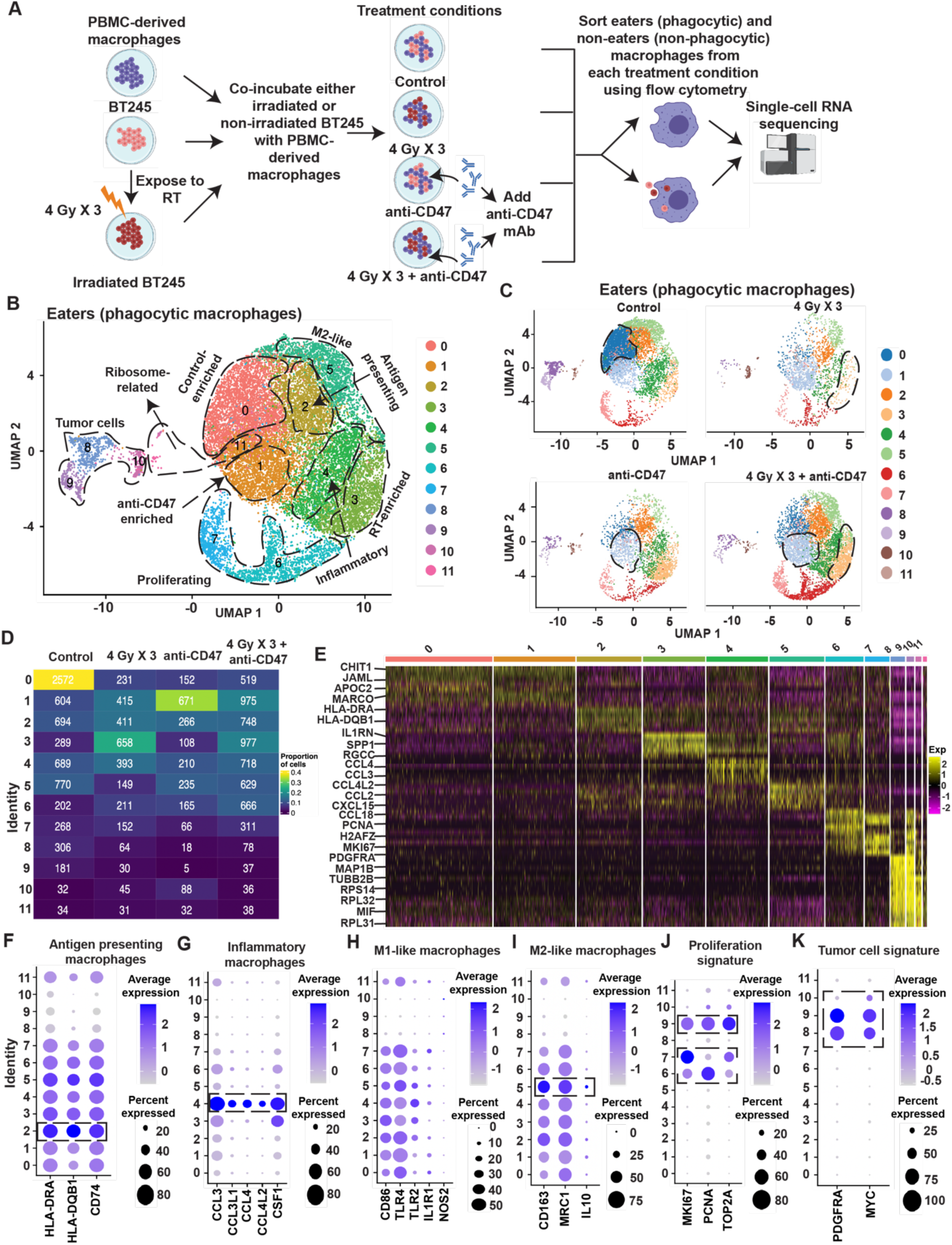
Single-cell RNA-sequencing reveals the enrichment of distinct macrophage subsets following co-culture with DMG cells pretreated with either Control, RT, anti-CD47 therapy, or combination of RT and anti-CD47 therapy. **(A)** Schematic diagram of the workflow used for *in vitro* phagocytosis and single-cell RNA-sequencing. Briefly, BT245 cells were irradiated (4Gy X 3) treated for three consecutive days and treated with either PBS or anti-CD47 mAb for 30 minutes at 37°C and co-cultured with PBMC-derived macrophages for 24 hours. BT245 cells not exposed to radiation and treated with PBS served as controls. Macrophages that either phagocytose (“eaters”) or don’t phagocytose (“non-eaters”) tumor cells were sorted using flow cytometry and subjected to single-cell RNA-sequencing. (**B)** UMAP projection displaying 11 distinct cell clusters from the eater’s cohort. Each dotted line and arrow indicate the identity of that specific cell cluster. **(C-D)** UMAP plots and proportion of cells in each cluster from 4 treatment groups: control, 4 Gy X 3, anti-CD47, or 4 Gy X 3 + anti-CD47. Dotted lines indicate the expansion/enrichment of distinct macrophage cluster in that treatment condition. Note the expansion/enrichment of two distinct cell clusters (1 and 3) in the combination treatment. (**E)** Heatmap of marker genes in each cell cluster. Representative genes with higher gene expression for each cluster are outlined on the left. Bubble plots demonstrating expression of marker genes associated with antigen presenting **(F)**, inflammatory **(G)**, M1-like (**H),** M2-like **(I),** proliferation **(J),** and tumor cell signature **(K)** by the various cell clusters. The dotted box indicates the cell clusters with higher average marker gene expression. The size of the bubble dot is proportional to the percentage of cells in a cluster expressing the marker gene and the color intensity is proportional to average scaled marker gene expression within a cluster.

Intriguingly, our analysis revealed that certain clusters exhibited striking preferential enrichment depending on the treatment condition of DMG cells, highlighting previously unappreciated immune responses induced by combination therapy.C0 was enriched in the control, C1 was enriched in the anti-CD47 treatment, and C3 was enriched in the RT treatment. There was an enrichment of both C1 and C3 following RT plus anti-CD47 treatment (**Figures 3C and 3D**). C2 was characterized by the expression of the antigen-presenting genes (HLA-DRA, HLA-DQB1, and CD74) (**Figures 3B, 3E and 3F**); C4 expressed inflammatory genes (CCL3, CCL3L1, CCL4, CCL4L2, and CSF1) (**Figures 3B, 3E and 3G**); C5 expressed anti-inflammatory M2-like signature genes (CD163, MRC1, IL-10) (**Figures 3B, 3E and 3I**). Interestingly, all the clusters expressed M1-like signature genes (CD86, TLR4, TLR2, IL1R1, NOS2) (**Figures 3B, 3E, and 3H**). C6 and C7 had higher expression of proliferation-related genes (KI67, PCNA, and TOP2A) (**Figures 3B, 3E and 3J**); C8, C9, and C10 expressed tumor-associated genes (PDGFRA and MYC) (**Figures 3B, 3E and 3K**). Tumor cells were identified by the exogenous expression of GFP, blasticidin, and Luciferase (Luc2) (transduced) (**Figures S5A, S5C, and S5E**). Lastly, C11 had upregulation of Ribosomal genes (RPS and RPL genes) (**Figures 3B, 3E and Table S3**).

We next performed QC and filtering of the scRNA-seq data obtained from the “non-eaters” (**Figure 3A**). In total, there were 6252 cells from the control, 4537 cells from the anti-CD47 treatment, 3541 cells from the RT, and 2060 cells from the RT plus anti-CD47 treatment (**Figure S6C**). Dimension reduction and clustering of the cells based on their gene expression profiles identified 13 different clusters (**Figure S6A**). Seven major macrophage clusters were identified and similar to the eaters, we saw a preferential enrichment of certain macrophage clusters depending on the treatment condition. For instance, C0 was enriched in the control, C1 was enriched in the RT treatment, and C2 was enriched in the control treatment, and C4 was enriched in the anti-CD47 treatment. Importantly, there was enrichment of both C1 and C4 following RT plus anti-CD47 treatment (**Figures S6B and S6C**). C0 and C5 expressed inflammatory genes (CCL3, CCL3L1, CCL4, CCl4L2, and CSF1) (**Figures S6A, S6D, and S6E)**; C3 expressed signature genes associated with lipid metabolism (CEBPA, APOC1, CYP27A1, GM2A, PLIN2, and ASAH1) (**Figures S6A, S6D, and S6E)**; C6, C9 and C10 expressed high levels of MKI67, PCNA, and TOP2A and were identified as proliferating cells **(Figures S6A, S6D, and S6G)**; C7, C10 and C13 expressed T cell signature genes (CD247, CD3D, CD3E, and CD3G) (**Figures S6A, S6D, and S6I)**; C8 had high expression of ribosomal biogenesis genes (FAU, RPS, and RPL proteins) (**Figures S6 A, S6D, S6H, and Table S4)**. Surprisingly, the cells in clusters 11 and 12 expressed tumor-associated genes such as PDGFRA, MYC, GFP, Luc2, and blasticidin (**Figures S5B, S5D, S5F, S6A, S6D, and S6J)**. This could be due to the carry-over of tumor cells from “eater” samples at the time of flow sorting or a subset of macrophages that had “eaten” tumor cells earlier is not double-positive for CD45 and Calcein-AM at the time of flow sorting (**Figure S4**).

### Distinct Macrophage Subsets Exhibit Treatment-Specific Transcriptional and Functional Adaptations in Response to DMG Cell Treatment-Induced Phagocytosis

To identify the genes and pathways that are altered in “eaters,” specifically in control-enriched (C0), anti-CD47 enriched (C1) and RT-enriched (C3) macrophages, we first determined the marker genes that were differentially expressed and distinguished control-enriched from anti-CD47 enriched and RT-enriched using the “FindMarkers” function in Seurat^32, 33^. We found control-enriched expressed higher levels of NUPR1, LYZ, and JAML and lower levels of IL1RN, SPP1, FABP4, TM4SF19, and RGCC (**Figure 4A and Table S5**). anti-CD47 enriched when compared to control-enriched and RT-enriched expressed higher levels of BLVRB, CHIT1, and HAMP and lower levels of IL1RN, SPP1, and ALCAM (**Figure 4C and Table S6).** RT-enriched compared to anti-CD47 enriched and control-enriched expressed higher levels of ILIRN, SPP1, FABP4, RGCC, and ALCAM and lower levels of CHIT1, ALOX5AP, GLUL, CD52, and PTGDS (**Figure 4E and Table S7**). Strikingly, genes upregulated in RT-enriched are downregulated in anti-CD47 enriched (**Figures 4A and 4E**). Next, to determine the pathways that are altered in control-enriched compared to anti-CD47 enriched and RT-enriched, we performed gene set enrichment analysis using clusterProfiler (an R package designed to perform over-representation analysis), enrichGO, and David bioinformatic analysis on the significant marker genes (p value adj < 0.01 and average Log_2_ FC > 0.25). Biological process in GSEA and Kyoto encyclopedia of genes and genomes (KEGG) pathway from DAVID identified lymphocyte proliferation, phagosome, endosome, and antigen processing and presentation via MHC class II genes are upregulated in control-enriched macrophages (**Figures 4B, S8A, and S8B**). Similar pathway analysis on differentially expressed significant marker genes in anti-CD47 enriched showed that ATP metabolic process, oxidative phosphorylation, electron transport chain, NADH dehydrogenase, mitochondrial ATP synthesis, reactive oxygen species, and antigen processing and presentation pathways via MHC class I are upregulated in anti-CD47 mAb treatment-enriched macrophages (**Figures 4D, S9A, and S9B**). In contrast, response to interferon-gamma, ERK signaling, leukocyte cell-cell adhesion, focal adhesion and monocyte chemotaxis pathways are upregulated in radiation-enriched macrophages (**Figures 4E, S10A and S10B**).

**Figure 4:**
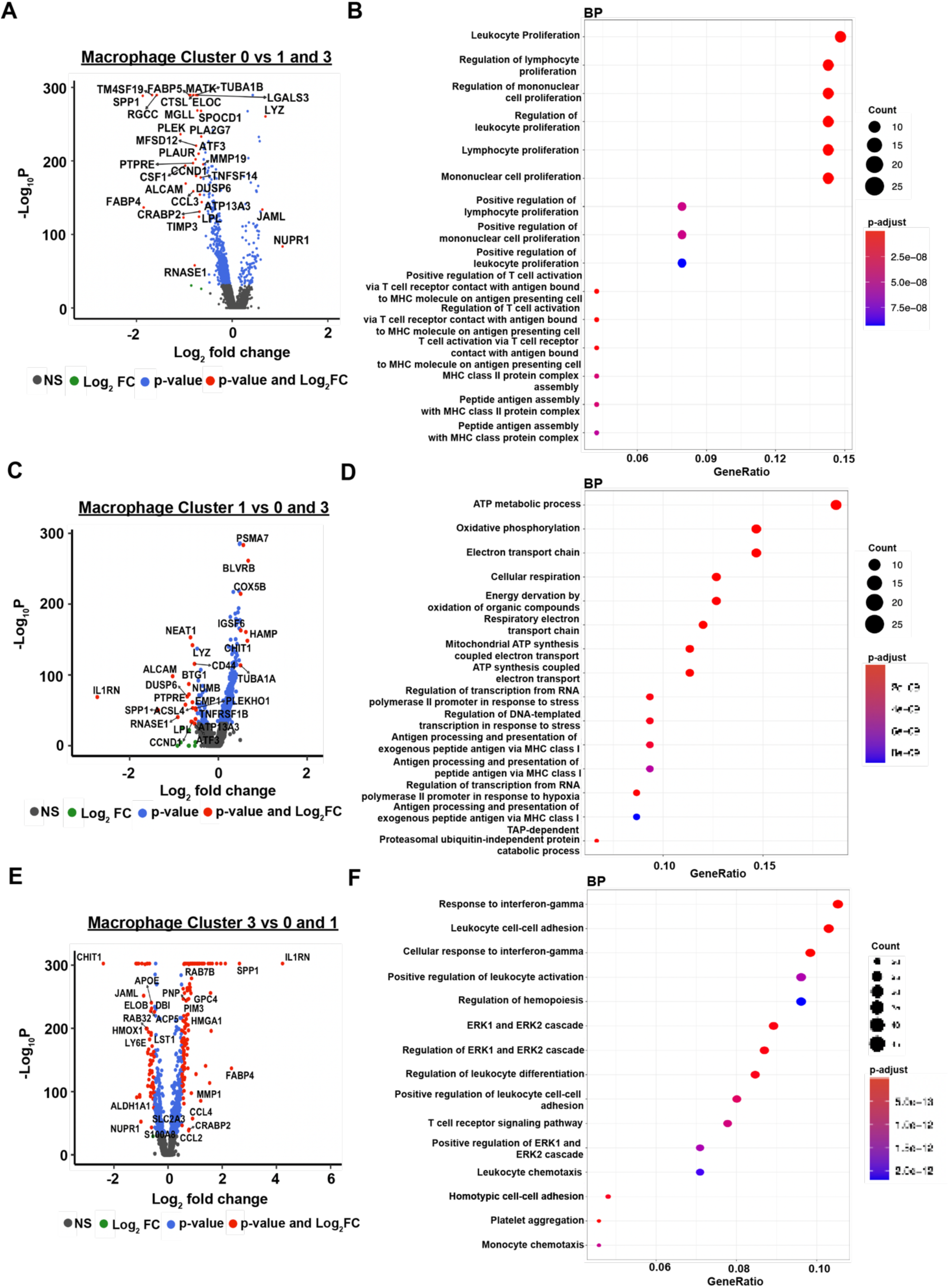
Characterization of macrophages that are enriched following phagocytosis of diffuse midline glioma cells pretreated with Control, anti-CD47 therapy or RT. **(A)** Volcano plot showing the top differentially upregulated and downregulated genes in the macrophages that are enriched after co-culture with control treated BT245 cells (cell cluster 0) compared to macrophages that were enriched after co-culture with either anti-CD47 (cell cluster 1) or RT (cell cluster 3) treated BT245 cells. (**B**) Gene Ontology enrichment analysis of biological process for significantly upregulated genes between control-enriched macrophages vs anti-CD47- and RT-enriched macrophages. Note only the top 15 biological processes are shown. (**C)** Volcano plot showing the top differentially upregulated and downregulated genes in the macrophages that are enriched after co-culture with anti-CD47 treated BT245 cells (cell cluster 1) compared to macrophages that were enriched after co-culture with either control (cell cluster 0) or RT (cell cluster 3) treated BT245 cells. (**D**) Gene Ontology enrichment analysis of biological process for significantly upregulated genes between anti-CD47-enriched macrophages vs Control- and RT- enriched macrophages. Note only the top 15 biological processes are shown. **(E)** Volcano plot showing the top differentially upregulated and downregulated genes in the macrophages that are enriched after co-culture with RT treated BT245 cells (cell cluster 3) compared to macrophages that were enriched after co-culture with either control (cell cluster 0) or anti-CD47 (cell cluster 1) treated BT245 cells. (**F)** Gene Ontology enrichment analysis of biological process for significantly upregulated genes between control-enriched macrophages vs anti-CD47- and RT- enriched macrophages. Note only the top 15 biological processes are shown.

To identify the genes and pathways altered in “non-eaters,” we performed an identical approach for “eaters.” In comparison to RT-enriched (C1) and antiCD47-enriched (C4) macrophages, control enriched macrophages (C2) expressed higher levels of PTGDS, CHIT1, CLU, MDM2, and CYP1B1 and lower levels of IL1RN, TM4SF19, RGCC, FABP4 (**Figure S7A and Table S8**). RT-enriched compared to control enriched macrophages and antiCD47-enriched expresses higher levels of MARCO, NUPR1, C1QA, CD14, and MARCKS and lower levels of IL1RN, RGCC, TM4SF19, CRABP2, and SPP1 (**Figure S7C and Table S9**). antiCD47-enriched compared to RT-enriched and control enriched macrophages expressed higher levels of IL1RN, RGCC, TM4SF19, SPP1, CRABP2 and lower levels of CD14, MARCKS, CTSC, NUPR1, and HLA-DMB (**Figure S7E and Table S10**). The pathway analysis on significant (p value adj < 0.01 and average Log_2_ FC > 0.25) marker genes in Control-enriched “non-eaters” revealed the upregulation of apoptotic signaling, p53 signaling, DNA damage checkpoint and oxidative phosphorylation (**Figures S7B, S11A, and S11B**). Upregulation of antigen processing and presentation, T cell receptor and CD4 receptor binding genes were observed in radiation-enriched “non-eaters” (**Figures S7D, S12A and S12B**). Lastly, upregulation of purine ribonucleotide metabolic process, ATPase and GTPase activity genes are upregulated in anti-CD47-enriched “non-eaters” (**Figures S9F, S13A, and S13B**).

Our analysis revealed distinct transcriptional adaptations in macrophages depending on their exposure to tumor cells pre-treated with either radiation or anti-CD47 therapy. Radiation-enriched macrophages exhibited upregulation of interferon-gamma response and ERK signaling, which may be attributed to radiation-induced DNA damage and immunogenic cell death (ICD). ICD leads to the release of damage-associated molecular patterns (DAMPs), including cytosolic DNA and ATP, which activate the STING pathway, subsequently inducing type I interferons and pro-inflammatory cytokines. This response can further enhance ERK signaling, a key pathway involved in macrophage activation, cytokine production, and inflammatory responses. In contrast, macrophages exposed to anti-CD47-treated tumor cells demonstrated significant upregulation of oxidative phosphorylation, ATP synthesis, and NADH dehydrogenase activity, suggesting that active live-cell phagocytosis is a metabolically demanding process. Engulfment and degradation of tumor cells require high ATP turnover to fuel cytoskeletal rearrangements, phagosome acidification, and lysosomal processing, leading to increased mitochondrial respiration. Notably, in macrophages that failed to phagocytose tumor cells, we observed upregulation of p53 signaling and DNA damage checkpoint pathways, which may indicate a stress response associated with prolonged tumor cell interaction without successful engulfment. The inability to clear tumor cells might lead to persistent DNA damage accumulation or oxidative stress, triggering cell-cycle arrest and apoptotic priming via p53 activation. These findings suggest that successful phagocytosis reprograms macrophage metabolism toward an oxidative state, whereas ineffective phagocytosis engages stress and DNA repair mechanisms, potentially altering immune functionality in the tumor microenvironment.

### Combination of radiotherapy and anti-CD47 treatment reduced tumor growth and prolongs the survival of human and mouse DMG/DIPG

The promising results from the *in vitro* phagocytosis assay and scRNA-seq prompted us to test the efficacy of combining RT plus anti-CD47 *in vivo* models of DMG/DIPG. To this end, we implanted BT245-Luc2 and SU-DIPGXVII-Luc2 in NOD-SCID IL2R gamma^null^ (NSG) mice. The schema for tumor establishment and RT and anti-CD47 treatments for the two respective models are outlined in **Figures 5A and S14A.** The above-mentioned lines were inoculated into the mouse pons and bioluminescence (BLI) imaging was used to monitor tumor establishment and tumor growth. Once tumors were established, mice were randomized into four groups (Control, RT, anti-CD47, and RT plus anti-CD47) and subjected to indicated treatment regimens and doses as shown in **Figures 5A and S14A.** The combination treatment of radiotherapy followed by anti-CD47 treatment reduced tumor growth (**Figures 5B, 5C, S14B, and S14C).** The mice receiving monotherapy (fractionated RT or anti-CD47 treatment) did not show any effect on tumor growth (**Figures 6B, 6C, S14B, and S14C).** The survival analysis suggested that the combination treatment significantly improved the survival compared to monotherapy, with the median survival increasing from 46 to 58 days in the BT245-bearing mice and from 170 to 210 days in the SU-DIPG17-bearing mice (P < 0.005). Importantly, in the less aggressive, SU-DIPG17 model, we see a complete regression of the tumor in 3/7 mice(S14D). There was no increase in the median survival of the mice that received either RT or anti-CD47 in either of the models (**Figures 6D and S14D**). Lastly, we assessed the tumor-bearing brain tissue at endpoint for its macrophage surface marker profile. In BT245 tumor bearing mice, that received anti-CD47 were found to have an increased number of F4/80+ macrophages when compared to all other treatments (**Figure 5E**). However, the mice that received the combination treatment showed an increased percentage of M1-like and decreased percentage of M2-like macrophages compared to all other treatments (**Figures 5F and 5G**). In the SU-DIPG17 model, the mice that received the combination treatment were found to have an increased number of F4/80+ and M1-like and decreased percentage of M2-like macrophages compared to all other treatments (**Figures S14E, S14F, and S14G**).

**Figure 5:**
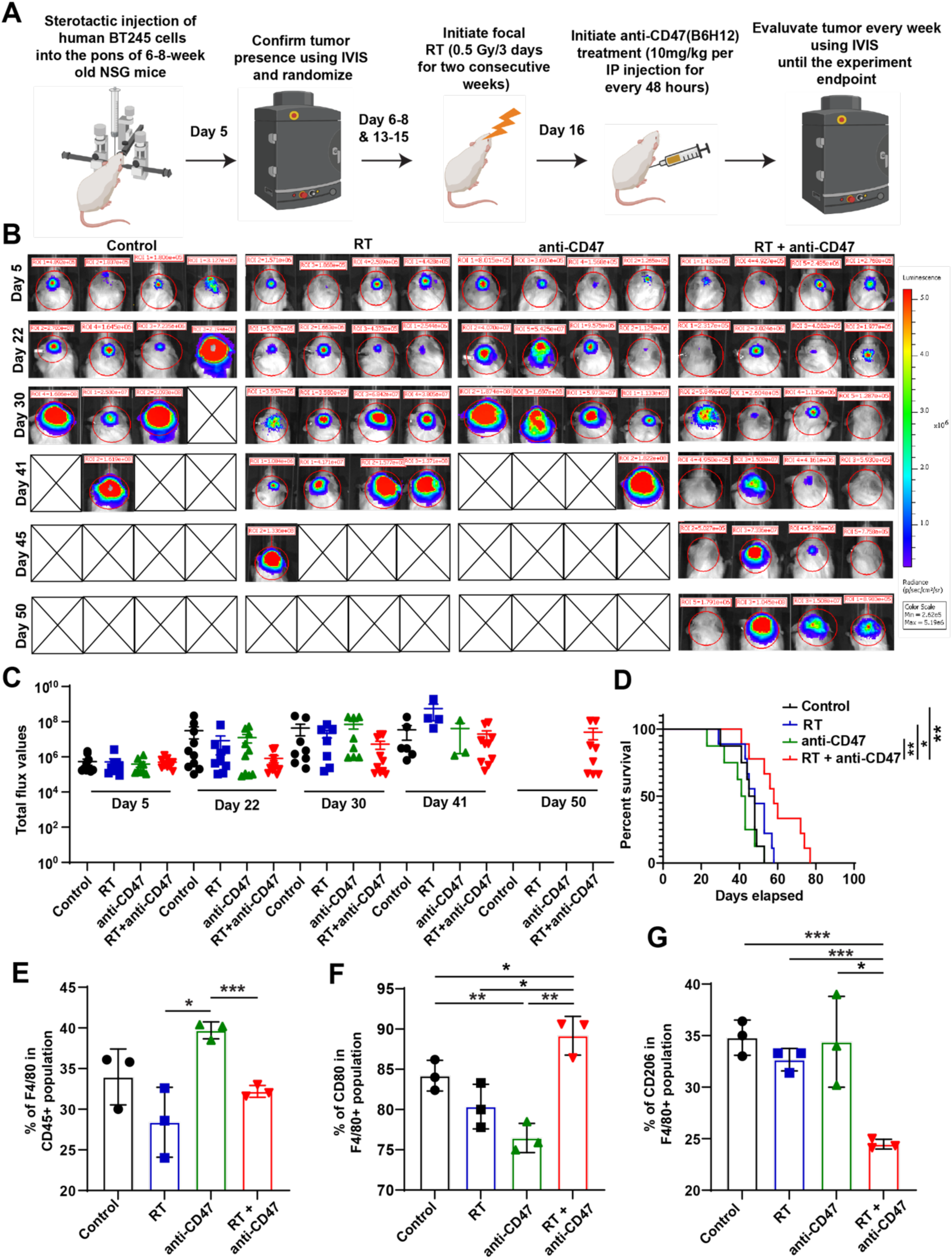
Combination of RT and anti-CD47 treatment reduces tumor burden and prolongs the survival of mice-bearing BT245 xenografts compared to monotherapy. **(A)** Schematic diagram showing the experimental treatment plan followed**. (B)** Representative bioluminescence images of the mice-bearing Luc2-expressing BT245 cells before, after, and during the specified treatments. The scale bar adjacent to the image displays bioluminescence counts (photons/second/cm^2^/steradian). **(C)** Quantification of total IVIS flux values over time course. **(D)** Kaplan-Meier survival analysis of BT245 xenografts with indicated treatments, control, n = 10; RT, n= 9; anti-CD47, n=10; and RT + anti-CD47, n=10. The log-rank test was used to calculate statistical significance. *p < 0.05, **p < 0.01. **(E-G)** Bar graphs demonstrating the relative percentages of F4/80^+^, CD80^+^ (M1-like) and CD206^+^ (M2-like) tumor-associated macrophages in control, RT, anti-CD47, or RT + anti-CD47 treated mice-bearing BT245 xenografts. Data shown are obtained from n=3 mice for each group and are represented as mean ± SD. Unpaired student’s t-test: *p < 0.05, **p < 0.01 and ***p < 0.0001.

**Figure 6:**
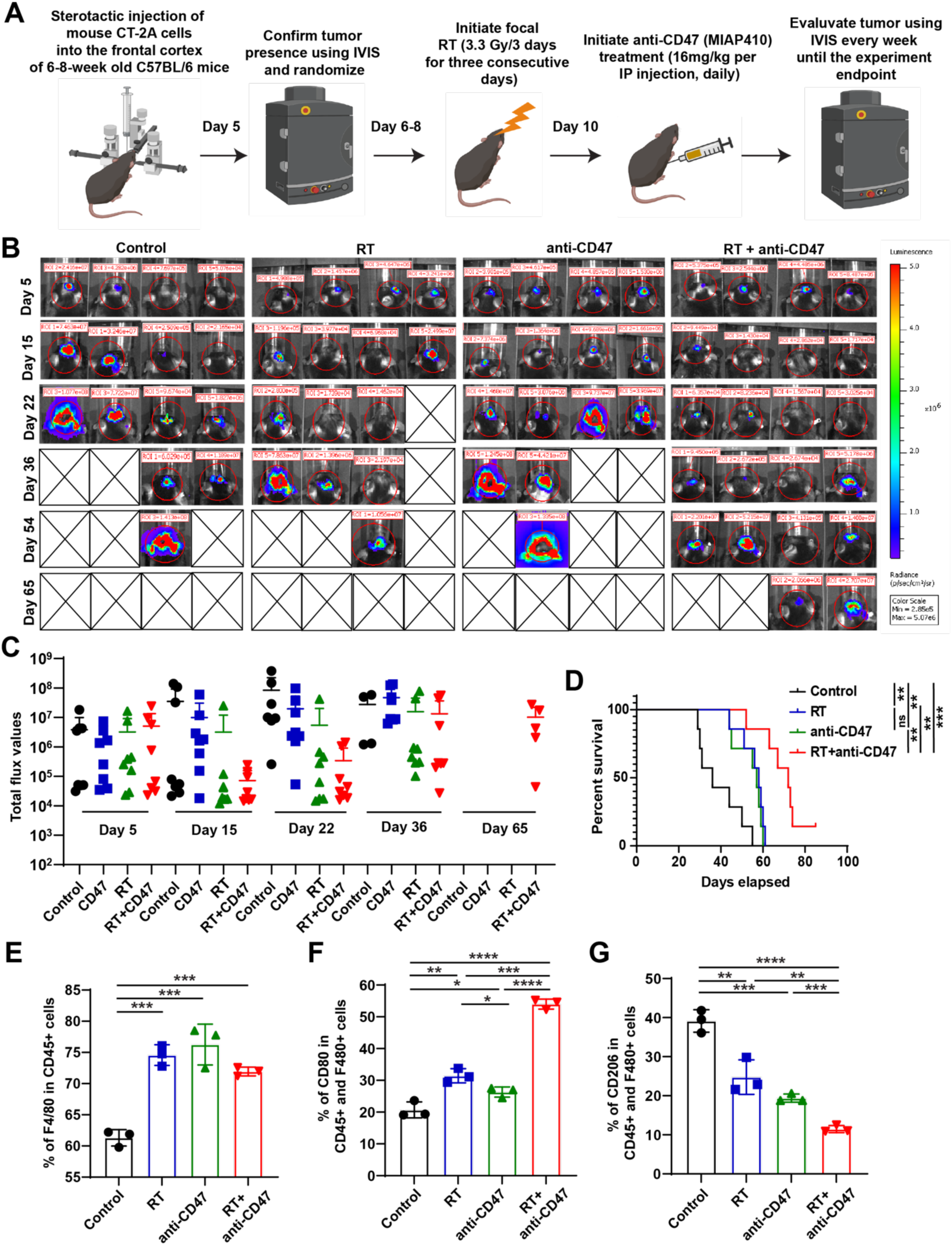
Combination of RT and anti-CD47 treatment reduces tumor burden and prolongs the survival of mice-bearing CT-2A xenografts compared to monotherapy. **(A)** Schematic diagram showing the experimental treatment plan followed**. (B)** Representative bioluminescence images of the mice-bearing Luc2-expressing CT-2A cells before, after, and during the specified treatments. The scale bar adjacent to the image displays bioluminescence counts (photons/second/cm^2^/steradian). **(C)** Quantification of total IVIS flux values over time course. **(D)** Kaplan-Meier survival analysis of CT-2A xenografts with indicated treatments, control, n = 8; RT, n= 8; anti-CD47, n = 8; and RT + anti-CD47, n = 8. The log-rank test was used to calculate statistical significance. *p < 0.05, **p < 0.01, ***p < 0.001. ns, not significant. **(E-G)** Bar graphs demonstrating the relative percentages of F4/80^+^, CD80^+^ (M1-like) and CD206^+^ (M2-like) tumor-associated macrophages in control, RT, anti-CD47, or RT + anti-CD47 treated mice-bearing CT-2A xenografts. Data shown are obtained from n = 3 mice for each group and are represented as mean ± SD. Unpaired student’s t-test: *p < 0.05, **p < 0.01 and ***p < 0.0001.

To determine the safety and efficacy of combining RT and anti-CD47 treatment in a syngeneic immune-competent mouse model we stereotactically implanted a GEMM derived syngeneic mouse pHGG cell line, PKC-HA into the pons of 6-8-week-old C57BL/6 mouse. The schema for tumor establishment and RT and anti-CD47 treatments for the two respective models are outlined in **Figure S15A**. Once tumors were established, mice were randomized into four groups (Control, RT, anti-CD47, and RT plus anti-CD47) and subjected to indicated treatment regimens and doses as shown in **Figure S15A.** In line with the findings from patient-derived xenograft (PDX) models, combining radiotherapy and anti-CD47 treatment reduced tumor growth compared to monotherapy (**Figures S15B and S15C)**. Additionally, the survival analysis suggested that the combination treatment significantly improved the median survival compared to monotherapy. Importantly, complete tumor regression was observed in 6/10 mice treated with the combination treatment. RT and anti-CD47 treatment also significantly increased the median survival compared to Control (**Figure S15D**). Taken together, these results strongly support that combining RT and anti-CD47 treatment is highly effective, well-tolerated and prolongs survival in DMG/DIPG animal models.

### Combination of radiotherapy and anti-CD47 treatment reduced tumor growth and prolongs the survival in immunocompetent GBM mouse models

Given that pediatric and adult high-grade gliomas (HGG) have historically exhibited differential responses to immunotherapies—likely due to variations in immune cell infiltration, mutational burden, and myeloid composition—we sought to investigate whether the promising effects of combining RT and anti-CD47 therapy observed in our pediatric models could be translated into adult HGG. Previous studies have highlighted the relative paucity of T-cell infiltration in pediatric HGG compared to their adult counterparts, which may influence the efficacy of myeloid-targeted therapies. We have recently shown that combining single dose of RT with anti-CD47 treatment significantly increased the survival in PDX models of GBM^24^. Nevertheless, the safety and efficacy of combining fractionated RT with anti-CD47 in GBM mouse models with intact immune system was not investigated previously. To this end, we stereotactically implanted two well-characterized mouse glioma lines CT-2A and SB28 into the frontal cortex of 6-8-week-old C57BL/6 mice. After implantation with CT-2A and SB28 luciferase-expressing cells, the tumor engraftment was confirmed by BLI, and mice were randomized into four groups (Control, RT, anti-CD47, and RT plus anti-CD47) and subjected to indicated treatment regimens and doses as shown in **Figures 6A and S16A.** Similar to the DMG/DIPG models, the combination treatment of radiotherapy followed by anti-CD47 treatment reduced tumor growth compared to monotherapy (**Figures 6B, 6C, S15C and S15D).** Additionally, the survival analysis of CT-2A-bearing mice suggested that the combination treatment significantly improved the median survival compared to monotherapy, with the median survival increasing from 36 to 72 days. RT alone and anti-CD47 alone also significantly increased the median survival to 58 and 57 days from 36 days (**Figure 6D)**. In the more aggressive, SB28-bearing mice, the combination treatment increased the median to 35 days from 30 days. In contrast, either RT alone or anti-CD47 alone did not significantly increase the median survival in SB28-bearing mice (**Figure S15D**). Next, we assessed the tumor-bearing brain tissue at endpoint for its macrophage surface marker profile. In the CT-2A-bearing mice, anti-CD47 treatment increased the infiltration of F480+ and M1-like macrophages and decreased the infiltration of M2-like macrophages compared to other treatments (**Figures 6E-6G**). In the SB28-bearing mice, both anti-CD47 alone and RT plus anti-CD47 treatment increased the increased the infiltration of F480+ and M1-like macrophages and decreased the infiltration of M2-like macrophages compared to other treatments (**Figure S16E-16G)**. Collectively, these results strongly support that combining RT and anti-CD47 mAb treatment is highly effective, well-tolerated, and prolongs survival in the fully immunocompetent GBM mouse models.

### Distinct Macrophage Subpopulations are Shaped by Radiation and Anti-CD47 Therapy in Gliomas

scRNA-seq revealed the enrichment of distinct macrophage subsets in response to co-culture with either control, irradiated, or anti-CD47 treated DMG cells. To validate these findings *in vivo* and at the protein-level, we first determined the marker genes that are expressed highly on the cell surface of control-enriched (C0), RT-enriched (C3) and anti-CD47-enriched (C1) macrophages. Several genes were expressed at higher levels, however, in comparison to all the cell clusters, CLEC7A showed higher expression in control-enriched (**Figure 7A**), CD44 showed higher expression in RT-enriched (**Figure 7D**), CD63 showed higher expression in antiCD47-enriched (**Figure 7G**). Next, to determine if these cell surface marker genes are enriched *in vivo*, we performed high dimensional flow cytometry (**Figure S17**) on single-suspension cells harvested at endpoint from the brains of C57BL/6 mice-bearing either PKC-HA or CT-2A cells and treated with different conditions as shown **Figures 6A and S15A.** In agreement with the scRNA seq data, we found higher expression of CLEC7A on the surface of F4/80+ macrophages infiltrating the brain tumors of control-treated mice compared to either RT or anti-CD47 or RT plus anti-CD47 treated mice (**Figures 7B and 7C**). Similarly, CD44 was expressed highly on the surface of F4/80+ macrophages infiltrating the brain tumors of RT and RT plus anti-CD47 treated mice compared to either control or anti-CD47 treated mice (**Figures 7E and 7F**). Finally, CD63 was expressed highly on the surface of F4/80+ macrophages infiltrating the brain tumors of anti-CD47 and RT plus anti-CD47 treated mice compared to either control or RT treated mice. In conclusion, these results validate the scRNA-seq data and confirm the existence as well as the enrichment of distinct macrophage subpopulations following treatment of tumor-bearing glioma mice with either control, RT, or anti-CD47 therapy. Strikingly, combination of RT and anti-CD47 treatment results in the enrichment of macrophage subpopulations that were enriched in either RT alone or anti-CD47 alone treated mice.

**Figure 7:**
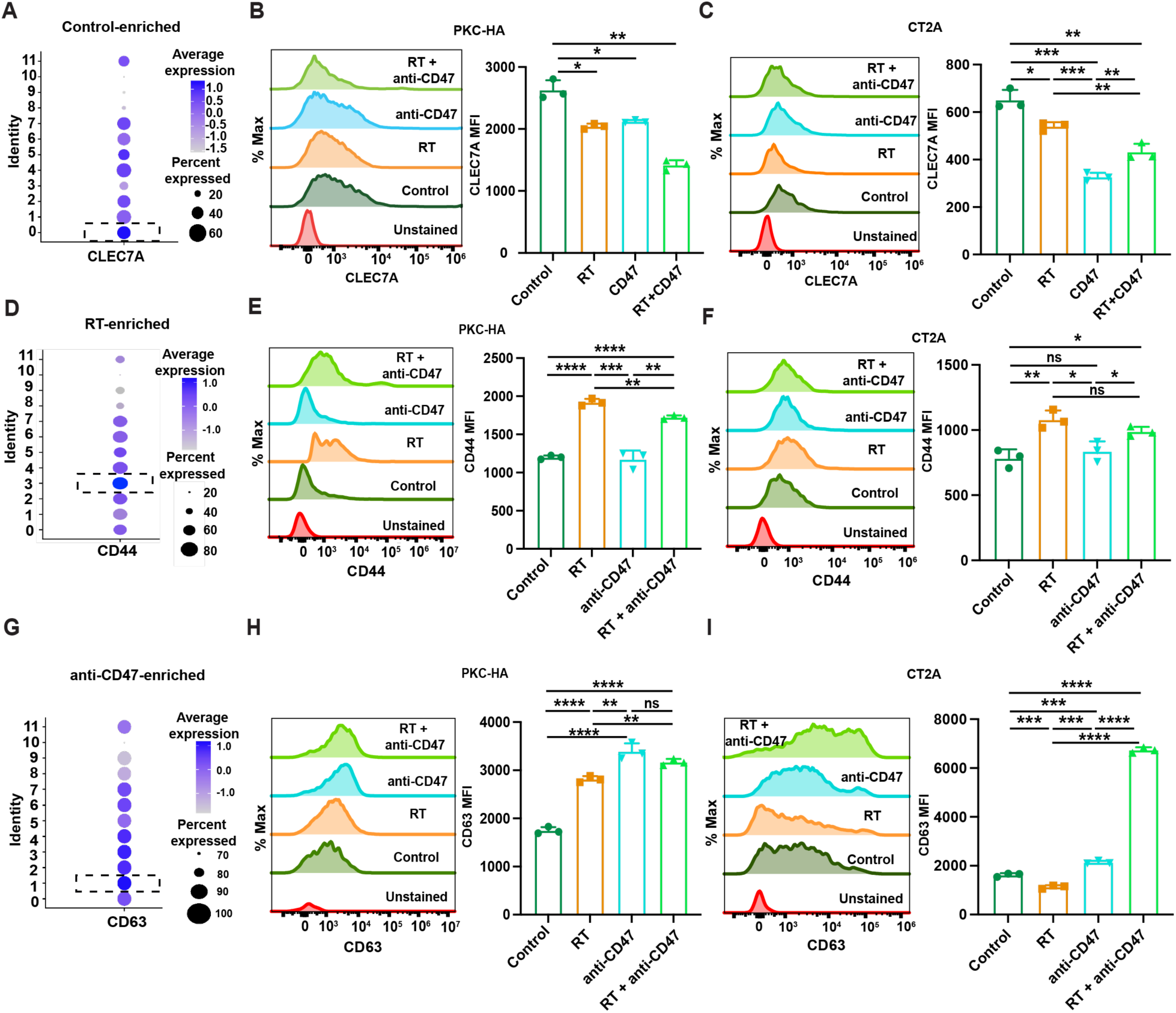
Validation of marker genes identified from single-cell RNA-sequencing using tumor-associated macrophages obtained from mouse DMG and glioma xenografts treated with either Control, RT, anti-CD47, or RT + anti-CD47. **(A)** Dot plot indicates the average expression of CLEC7A, CD44 (D), and (G) for each cell cluster identified from scRNA-seq. **(B-C)** Representative overlay histograms and median fluorescence intensity (MFI) values of CLEC7A, CD44 (**E-F),** and CD63 **(H-I)**, expression in gliomas isolated from mice treated with either PBS (control), RT, anti-CD47 therapy, or RT with anti-CD47 combination therapy. Data shown are obtained from n=3 mice for each group and are represented as mean ± SD. ns, not significant; Unpaired student’s t-test: *p < 0.05, **p < 0.01 and ***p < 0.0001.

## Discussion

This study reveals that macrophage fate is intricately linked to what they phagocytose, challenging the traditional notion of phagocytosis as a uniform process. Instead, phagocytosis emerges as a dynamic and multifaceted event that shapes macrophage transcriptional, metabolic, and functional states in response to the specific stimuli encountered in the tumor microenvironment. Radiotherapy (RT) and anti-CD47 monoclonal antibody (mAb) treatment significantly alter the immunogenic landscape of tumor glioma cells and consequently, the identity and function of macrophages that interact with these cells. These findings highlight the importance of considering phagocytosis not only as an endpoint of immune activation but also as a driver of macrophage specialization and plasticity.

The induction of immunogenic cell death (ICD) by RT, characterized by the dose-dependent upregulation of damage-associated molecular patterns (DAMPs) such as phosphatidylserine (PS), calreticulin (CRT), heat shock proteins (HSP70 and HSP90), and the extracellular release of high mobility group protein 1 (HMGB1), profoundly influences macrophage function. By exposing or releasing these immunogenic signals, RT primes tumor cells for recognition and uptake by macrophages. However, the macrophage response to phagocytosis of these irradiated cells is not homogeneous. Instead, RT-induced tumor cell death shapes a distinct macrophage phenotype enriched in inflammatory and interferon-gamma-responsive pathways. These macrophages exhibit transcriptional upregulation of ERK signaling and antigen presentation pathways, suggesting that the nature of what macrophages “eat” directly programs their downstream functions and immune-stimulatory capacity.

Similarly, anti-CD47 mAb treatment, which blocks the “don’t eat me” signal, enhances macrophage-mediated phagocytosis but elicits a different transcriptional and metabolic response. Macrophages that engulf tumor cells treated with anti-CD47 mAb upregulate metabolic pathways, including oxidative phosphorylation and ATP synthesis, and exhibit enhanced antigen presentation through MHC class I molecules. This metabolic reprogramming underscores that phagocytosis is not merely an end-point event but a process that initiates distinct macrophage functional programs, driven by the immunologic and molecular properties of the ingested material.

The combination of RT and anti-CD47 therapy revealed a remarkable synergy in macrophage-mediated tumor clearance, with macrophages adopting a hybrid phenotype that incorporates features of both RT- and anti-CD47-enriched subsets. Single-cell RNA sequencing (scRNA-seq) revealed that macrophages in the combination-treated groups exhibited transcriptional signatures of metabolic activation, inflammatory signaling, and enhanced antigen presentation. These hybrid macrophages likely represent an optimal immune phenotype for anti-tumor activity, integrating the immune priming effects of RT with the enhanced phagocytic capacity enabled by anti-CD47 treatment. This finding reinforces the idea that macrophage fate is not only shaped by the process of phagocytosis but is also highly context-dependent, influenced by the molecular properties of the material being internalized and the broader microenvironmental signals.

The in vivo data further supports the idea that macrophage fate is determined by the immunogenicity of the material they phagocytose. Tumor-associated macrophages (TAMs) from mice treated with RT and anti-CD47 therapy showed a marked shift in polarization toward an M1-like, pro-inflammatory phenotype, with reduced representation of M2-like, pro-tumorigenic macrophages. Importantly, the enhanced anti-tumor macrophage activity was associated with a significant reduction in tumor growth and, in some models, complete tumor regression. These novel findings highlight the plasticity of macrophages and their ability to adopt distinct functional states based on the nature of the tumor cells they ingest.

The study’s findings challenge the conventional view of phagocytosis as a singular process and instead present it as a critical determinant of macrophage identity and function. Macrophages are not passive scavengers but active participants in the immune response, with their fate and activity directly influenced by the molecular composition of what they phagocytose. For instance, macrophages that engulf irradiated tumor cells rich in DAMPs adopt an inflammatory, immune-activating phenotype, while those that internalize CD47-inhibited cells reprogram metabolically to optimize antigen processing and presentation. This plasticity underscores the importance of understanding phagocytosis as a dynamic process that shapes immune responses in diverse ways, depending on the context.

The implications of these findings extend beyond gliomas, as the principles of macrophage plasticity and the dynamic nature of phagocytosis are likely relevant to other cancers and diseases involving macrophage activity. For example, macrophage phenotypes in infectious diseases, autoimmune conditions, and tissue repair may also be determined by the specific material they phagocytose. Understanding these context-dependent interactions could inform strategies to reprogram macrophages in various disease settings by altering the nature of the material they encounter.

Future research should aim to dissect the molecular mechanisms underlying the macrophage specialization observed in this study. For example, how do specific DAMPs or CD47 blockade signals program distinct transcriptional and metabolic pathways in macrophages? Additionally, the interplay between phagocytosis and other macrophage functions, such as cytokine production, antigen presentation, and tissue remodeling, warrants further investigation. Identifying the factors that influence macrophage fate could lead to novel therapeutic strategies to modulate macrophage activity in cancer and other diseases.

In conclusion, this study highlights that macrophage fate is intimately linked to the molecular properties of what they phagocytose. Phagocytosis is not a singular, uniform process but a dynamic and context-dependent event that drives macrophage specialization and plasticity. By demonstrating that RT and anti-CD47 therapy shape distinct macrophage phenotypes through their effects on tumor immunogenicity, this study provides a framework for understanding how to harness and reprogram macrophages activity for therapeutic benefit. These findings underscore the potential of targeting macrophage plasticity as a strategy to enhance anti-tumor immunity and improve outcomes in malignant gliomas and other diseases.

## Materials and Methods

### Cell lines and cell cultures

DIPG/DMG cells were maintained as previously described^12^. Briefly, SU-DIPXVII and SU-DIPGXXV cells were provided by Dr. Michelle Monje (Stanford University, California) and cultured in tumor stem media (TSM) consisting of Neurobasal (-A), human basic fibroblast growth factor (FGFb; 20ng/ml; Shenandoah Biotech), human epidermal growth factor (EGF; 20ng/ml; Shenandoah Biotech), human PDGF-AA (20ng/ml; Shenandoah Biotech), PDGF-BB (20ng/ml; Shenandoah Biotech), and heparin sulfate (10ng/ml) (Sigma). BT245 cells were provided by Dr. Keith Ligon (Dana-Farber Cancer Institute, Boston) and maintained in human NeuroCult NS-A media (StemCell Technologies, Inc) supplemented with penicillin/streptomycin (100x), heparin (2 mg/ml), human epidermal growth factor (EGF; 20ng/ml), and human basic fibroblast growth factor (FGFb; 10 ng/ml). Mouse pediatric high-grade glioma cell lines, PKC-HA and PHC-HA were provided by Dr. Oren Becher and grown in mouse NeuroCult NS-A media (StemCell Technologies) supplemented with penicillin/streptomycin, heparin (2 mg/ml, StemCell Technologies), human epidermal growth factor (EGF; 10ng/ml), and human basic fibroblast growth factor (FGFb; 20 ng/ml). Mouse glioma cell line, SB28 cells were provided by Dr. Hideho Okada (University of California San Francisco) and grown in RPMI media 1640 (Gibco) supplemented with 10% (v/v) heat inactivated fetal bovine serum (FBS), penicillin/streptomycin (100x), GlutaMax (100x), MEM non-essential amino acids (10 mM), HEPES buffer solution (1M), Sodium Pyruvate (100 mM), and 2-mercaptoethanol (0.05 mM). The CT-2A cell line was purchased from Millipore Sigma and maintained in RPMI media 1640 as described above. Cells were grown in either as tumor neurospheres (ultra-low attachment plates or flasks, Corning) or adherent monolayer conditions (CytoOne) as indicated. All patient-derived cell lines were authenticated using DNA fingerprinting (STR analysis) at the Molecular Biology service center (UCD-AMC). The cell lines were routinely tested for mycoplasma contamination using Venor^TM^ GeM Mycoplasma detection kit (Sigma-Aldrich, St. Louis, MO). The list of cell lines used in this study are shown in **Table S1**.

### Irradiation

#### *In vitro* and IC_50_ determination

Cell lines used for *in vitro* assays were cultured on Geltrex-coated 96-well plate or T-25 flasks for 24 hrs. Cells were subjected to irradiation (0 Gy X 3, 2 Gy X 3, or 4 Gy X 3) for 3 consecutive days using ^137^Cs irradiator. All irradiations were performed using a perpendicular 662 KeV g-photon beam with dose coverage to uniformly irradiate all samples at a dose rate of 1.09 Gy/minute and a source-to-surface distance of 30 cm. To determine the IC_50,_ the cells were irradiated as described above and on Day 5 after post-irradiation, cell viability was measured as the intracellular ATP content using the CellTiter-Glo Luminescent Cell Viability Assay (Promega), following the manufacturer’s instructions. IC_50_ was calculated using GraphPad prism software.

#### In vivo

NOD-scid IL2rgnull (NSG) mice implanted with BT-245 and SU-DIPGXVII cells received fractionated doses of 1 Gy per day for 3 consecutive days over 2 weeks (a total of 6 Gy). C57BL/6 mice implanted with PKC-HA, SB28, and CT-2A received 3.3 Gy per day for 3 consecutive days (total 10 Gy). Under isoflurane anesthesia, each mouse was positioned in the prone orientation and aligned to the isocenter in two orthogonal planes by fluoroscopy. Each side of the mouse brain received half of the dose, which was delivered in opposing, lateral beams. Dosimetric calculation was done using a monte-carlo stimulation in SmART-ATP (SmART scientific solutions B.V.) for the midbrain + pons + frontal lobe receiving the prescribed dose. Treatment was administered using a 225 KV photon beam with 0.3 mm Cu filtration through a circular 10-mm diameter collimator.

### Flow cytometry

Irradiated human DIPG/DMG and mouse glioma cell lines were dissociated by treatment with TrypLE and rinsed with cell staining buffer (BioLegend). Single cell suspensions of tumor cells were incubated with Truestain FC blocking solution (Biolegend) for 10 mins at 4°C. Next, the Zombie fixable aqua dye (Biolegend) diluted (1:1000) in PBS was added and samples were incubated for 20 minutes 4°C. Lastly, a cocktail of fluorophore-conjugated antibodies resuspended in 100 µl cell staining buffer (Biolegend) was added directly to each tube, and samples were incubated at 4°C for 30 min in the dark. All the antibodies used in this study are shown in **Table S2.** Flow cytometry was performed using CytoFLEX LX (Beckman Coulter). FlowJo v10.8 was used for data analysis.

### Elisa assay

BT-245, SU-DIPGXVII, SU-DIPGXXV, CT-2A and SB28 were exposed to increasing doses for RT (**Figure 1E and S1E**) for 3 consecutive days. 48 hours post the final RT treatment, HMGB1 concentration in the supernatants of irradiated human DMG/DIPG and mouse glioma cell lines were quantified by using human and mouse HMGB1 Elisa kit (Novus Biologicals) according to the manufacturer’s instructions.

### *In vitro* phagocytosis

In vitro phagocytosis assay with human and mouse macrophages were performed as described previously^12, 24, 34^. To obtain human monocytes, leukopak from healthy individuals were obtained from Children’s Hospital of Colorado Blood Conor Center. The blood was diluted 1:1 with PBS (Gibco) and PBMCs were separated on a Ficoll density gradient (GE healthcare). CD14+ monocytes were positively selected by using MojoSort Human CD14 selection kit (BioLegend), then plated at 1 × 10^6^/ml in 150 × 25 mm (Corning) non-treated sterile tissue culture plates in RPMI with 10% FBS, 1x penicillin/streptomycin, 200 mM glutamine, and 25 mM HEPES (all from Corning). To generate monocyte-derived macrophages, monocytes were treated with human recombinant M-CSF (50 ng/ml) (Pepro Tech) and human recombinant GM-CSF (10 ng/mL) (Pepro Tech). After 48 hours, non-adherent cells were removed and the cells were washed with either 1X PBS or 1X HBSS twice, and above-mentioned media with growth factors were added, and the plates were returned into the tissue culture incubator for an additional 5 days. Mouse macrophages are obtained from 7- to 10-week-old C57BL/6 mouse bone marrow. The mice were killed in a CO_2_ chamber and femur and tibiae were isolated. The bones were kept in ice-cold PBS and sterilized in 70% ethanol. By flushing them with mouse macrophage medium (RPMI with 10% FBS, 1x penicillin/streptomycin, 200 mM glutamine, and 25 mM HEPES), bone marrow cells were collected and plated at 1 × 10^6^/ml in 150 × 25 mm non-treated tissue culture plates in mouse macrophage medium containing mouse recombinant M-CSF (50 ng/ml) (Pepro Tech) and mouse recombinant GM-CSF (10 ng/mL) (Pepro Tech). The media was changed after 48 hrs and both human and mouse macrophages were harvested using TrypLE (Gibco) for experiments on days 8-9. Unless mentioned otherwise, at the end of differentiation protocol, dissociated macrophages and Mock (0 Gy) or irradiated human DIPG/DMG, and mouse glioma cell lines stained with CFSE (Biolegend) were co-cultured in macrophage medium as described above with or without prior addition of 50µg/ml anti-CD47 monoclonal antibody (H5F9-G4) at 37°C for 2 hours in ultra-low attachment round bottom plate (Corning) 96-well plates. The tumor cell to macrophage ratio was 2:1. Cells were analyzed with CytoFLEX LX (Beckman Coulter) using a high throughput autosampler. Gates were placed according to unstained and fluorescence-minus-one (FMO) controls. DAPI, Zombie Aqua and NIR fixable viability kit (Biolegend) were used to exclude dead cells. The flow cytometry gating strategy for determining the percentage of tumor cell phagocytosis by macrophages is shown in **S3**. Phagocytosis assays for each tumor type were performed in triplicates and repeated at least two times.

### RNA extraction from sorted cells, cDNA, and library preparation for single-cell RNA-seq

A live single cell suspension of “eating” (phagocytic) and “non-eating” (non-phagocytic) macrophages as shown in **Figure 3A** were flow sorted and single-cell library preparation and RNA-sequencing was performed at the University of Colorado Anschutz medical campus genomics and micro array core.

### Single-cell RNA-seq data analysis, clustering, and visualization

Raw sequencing reads were processed using Cell Ranger single-cell software suite (v 6.1.1) with default parameters. Reads were aligned to the human reference genome (GRCh38 V3.0.0) from the 10x Genomics website. The resulting count table was filtered for features expressed by at least five cells and cells with at least 500 detected features. Cells with a small number of gene counts (< 500) or high mitochondrial counts (mt-genes > 0.2) were filtered out. After removing unwanted cells, gene expression measurements for each cell were normalized using the “LogNormalize” function in Seurat (v 4.0.4)^35^. Differentially expressed genes were identified using the “FindAllMarkers” function in Seurat (v 4.0.4)^35^. Genes with fold change more than 1.5 and adjusted P value less than 0.05 were considered as differentially expressed genes. Cell clustering was performed using the “RunUMAP’’ and “FindClusters” function in Seurat with setting parameters dims=1:10 and resolution= 0.6 and visualized using Uniform Manifold Approximation and Projection (UMAP). Marker genes were used to assign identity to cell types.

### Pathway enrichment analysis

To identify differentially expressed molecular pathways in enriched macrophage subpopulation, DAVID bioinformatics resources (v 6.8)^36^ was performed on differentially expressed significant marker genes. Gene sets from Gene Ontology (GO) biological process^37^ and Kyoto Encyclopedia of Genes and Genomes (KEGG)^38, 39^ were used.

### Cell cycle scoring

Cell cycle phases of each cell were determined using the cell cycle scoring function in Seurat (v 4.0.4)^35^. The cell clusters that had high expression for KI67, PCNA, and TOP2A were considered proliferative.

### Orthotopic Xenograft models for malignant gliomas (Diffuse midline Glioma/Diffuse Intrinsic Pontine Glioma and Glioblastoma Multiforme)

*In vivo* mouse xenograft studies, including implantation, animal care, radiation, treatments, and euthanasia were performed under and approved by the University of Colorado Institutional Animal Care and Use Committee (IACUC) protocol #00777. Early passage BT-245 and SU-DIPG XVII cells that were transduced with Luc2-GFP are dissociated into single cells and orthotopically implanted in the pons of 8- to 10- week-old NSG mice. Briefly, mice were anesthetized and immobilized in a stereotactic frame. A midline incision was made on the skin to expose the scalp, and a microdrill was used to perform craniotomy 0.8 mm lateral to midline, 0.5 mm posterior to lambda, and 5.00 mm ventral to the surface of the skull. Next, a suspension of ∼1 × 10^5^ cells in 2 ml serum-free media were stereotactically injected at the rate of 500 nl/minute with a Hamilton syringe (ga26s/51mm/pst2). For the syngeneic model of DMG/DIPG, a suspension of ∼5 × 10^5^ PKC-HA cells were stereotactically injected into the pons of 6- to- 10-week-old C57BL/6. For the syngeneic model of GBM, ∼SB28 (5 × 10^4^) and GL261 (2.5 × 10^4^) cells were stereotactically injected at the rate of 500 nl/minute into the brain at a site of 2 mm lateral to bregma and 3mm ventral to the surface of the skull. After leaving the needle in place for 1 min, it was retracted at 3mm/min and the burr hole was closed with sterile bone wax (Ethicon, inc), the skin was sutured and placed in a cage atop a surgical heat pad until ambulatory. Tumor formation was monitored by bioluminescent imaging (BLI) using IVIS Xenogen 2500 imaging machine. After confirmation of intracranial tumor establishment with a total flux from BLI corresponding to ∼10^5^ to 10^6^ photons/second/cm^2^, animals were assigned into different treatment groups using www.randomizer.org. Following randomization, mice were injected with luciferin intraperitoneally and imaged for tumor progression once weekly until the end of the study. Body weight was measured once a week and mice were monitored daily and those reaching end point were euthanized according to IACUC protocols by CO_2_ asphyxiation, when they show signs of either neurologic deficit, failure to ambulate, body score less than 2, or weight loss greater than 15%.

### Antibody treatment

Mouse monoclonal anti-CD47 antibody (clone B6H12) was purchased from Bio X cell (West Lebanon, NH) and diluted in PBS (Phosphate Buffered Saline) before use. All the NSG mice implanted with human glioma cells started receiving intraperitoneal (IP) injections of 16 mg/kg of anti-human CD47 mAb on day 18 till the end of the study as depicted in **Figure 5A and S14A**. All the C57BL/6 mice with human glioma cells started receiving intraperitoneal (IP) injections of 32 mg/kg of anti-mouse CD47 ab (clone MIAP410) on day 18 till the end of the study as depicted in **Figure 6A, S15A, and S16A.**

### Tumor tissue dissociation

After resection of malignant gliomas from mice at endpoint, tumor samples were placed on ice and cut into small pieces and single-cell suspensions were made as described previously^12^. Briefly, tumors were enzymatically dissociated by collagenase IV (1mg/ml) (Worthington) in dissociation solution containing HBSS with calcium/magnesium (Gibco), non-essential amino acids (Gibco), sodium pyruvate (Gibco), sodium bicarbonate (Gibco), 25mM HEPES (Gibco), 1X Glutmax-1 (Gibco), antibiotic-antimycotic (Gibco), DNase (Worthington) at 37° C. The suspension was washed two times with HBSS and filtered using 70 mM and 40 mM filters, respectively. The cells were further treated with ACK/RBC lysis buffer (Gibco), washed once with HBSS, resuspended in Bambanker (Fisher Scientific), and stored in liquid nitrogen until use. For multicolor flow cytometry, cells were thawed from liquid nitrogen, blocked, and stained as described under flow cytometry section.

### Statistical analysis

All statistical analysis was performed using Graph pad prism V.9.2.0. Results were expressed as mean ± SD. One-way ANOVA followed by Bonferroni’s post-hoc comparison tests were used for three or four group comparisons and two-tailed unpaired Student’s t-tests were used for comparing two conditions. Animal survival curves were analyzed using Kaplan-Meier method and Grehan-Breslow-Wilcoxon tests, with groups compared by respective median survival or number of days taken to reach 50% morbidity. Differences were considered significant at P < 0.05 (*).

## Figures

**Figure 3A**, **Figure 6A, S14A, S15A, and S16A** were created using biorender.com.

## Data availability

The data generated in this study are available within the article and its Supplementary Data files. scRNA-sequencing data generated in this study are publicly available in Gene Expression Ombnibus (GEO) at GSE202174.

## Code availability

Custom scripts for analyzing single-cell RNA-sequencing data are available upon request.

## Materials and Correspondence

All request for materials and correspondence may be sent to

## Acknowledgements

We acknowledge the services provided by flow cytometry core and genomics core. The UCD cancer center core facilities are supported by NCI cancer center support grant no. P30-CA046934.

## Supplementary Figures

**Supplementary Figure 1:**
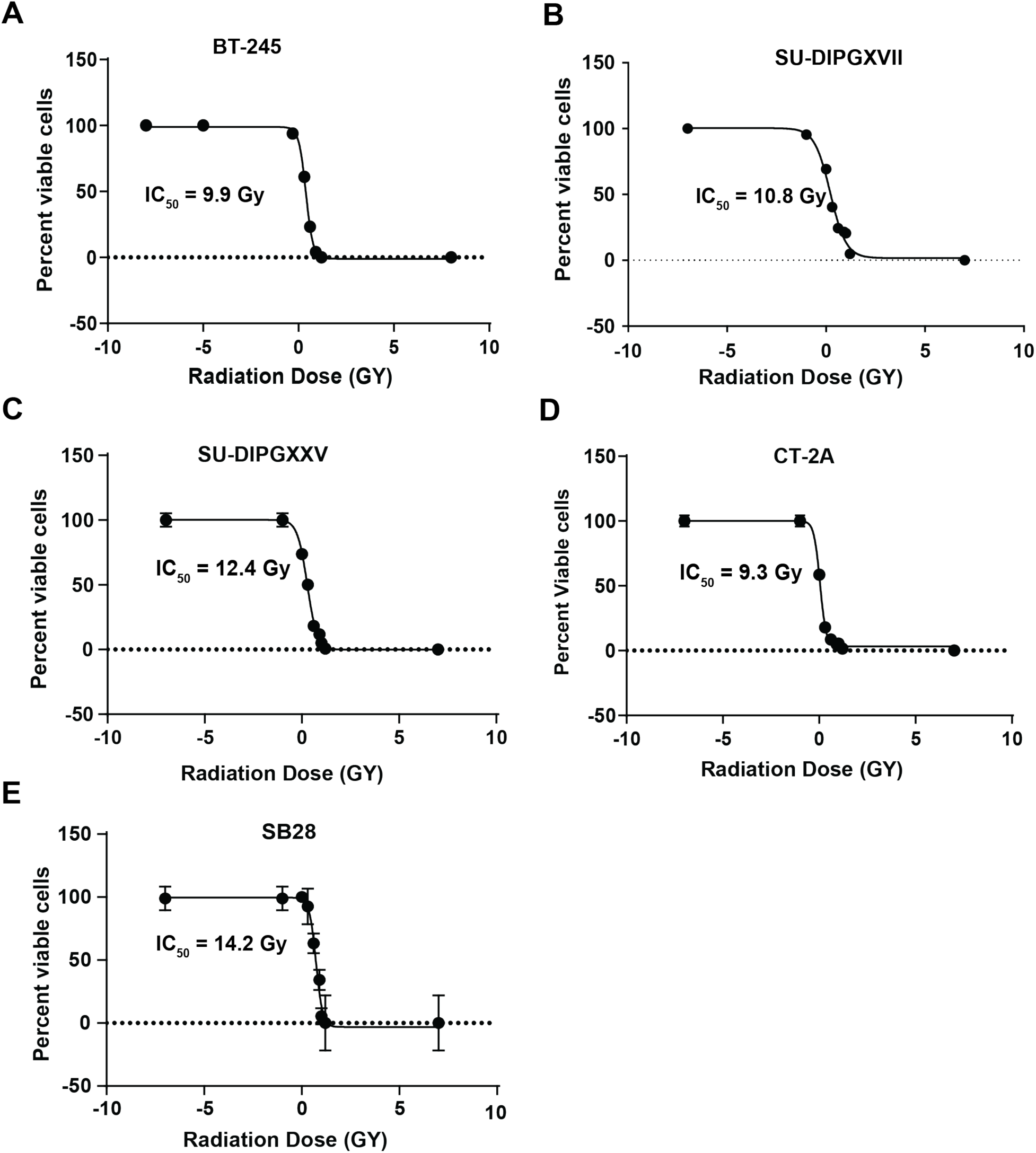
IC_50_ of human DMG and mouse glioma cell lines. (A-C) DMG cell lines and (E and F) mouse glioma cell lines were exposed to increasing doses of RT for three consecutive days and subsequently cell proliferation was measured on day 5 and IC_50_ of radiation was calculated.

**Supplementary Figure 2.**
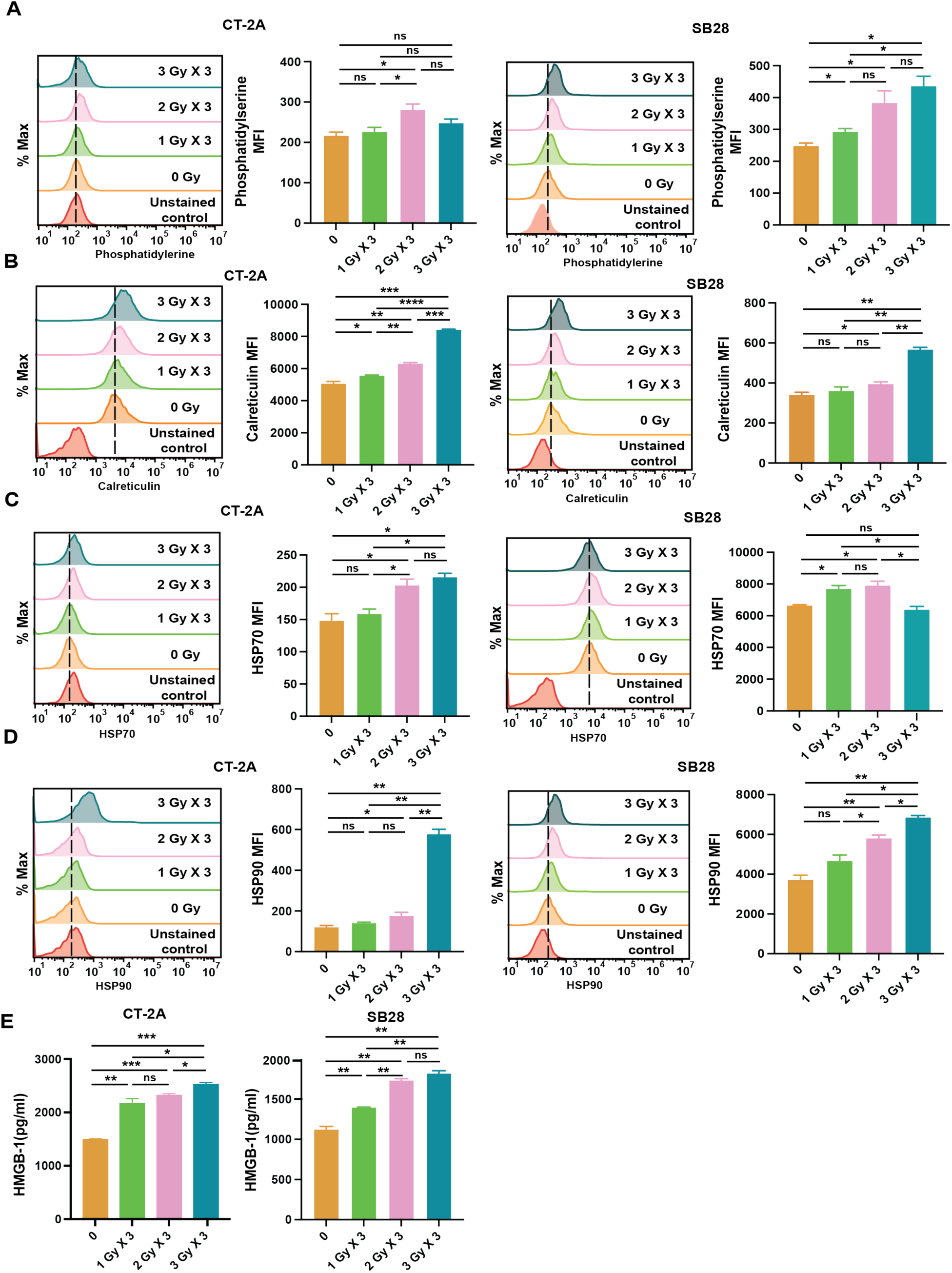
Radiation therapy induces the surface expression and release of damage-associated molecular patterns (DAMPs) in mouse glioma cell lines. Mouse glioma cell lines CT-2A and SB28 were exposed to increasing doses of RT for 3 consecutive days. **(A-D)** Representative overlay histograms and median fluorescence intensity (MFI) values displaying the expression levels of phosphatidylserine (PS), Calreticulin (CRT), Heat shock protein (HSP70), and HSP 90 on the surface of CT-2A and SB28 cells. Results are expressed as mean ± SD (n=3 technical replicates). ns, not significant. Unpaired student’s t-test: p < 0.05, **p < 0.01. **(E)** 24 hours post the final RT treatment, HMGB1 released in the supernatants of CT-2A and SB28 cells were quantified. Results are expressed as mean ± SD (n=3 technical replicates). ns, not significant; Unpaired student’s t-test: p < 0.05, **p < 0.01.

**Supplementary Figure 3.**
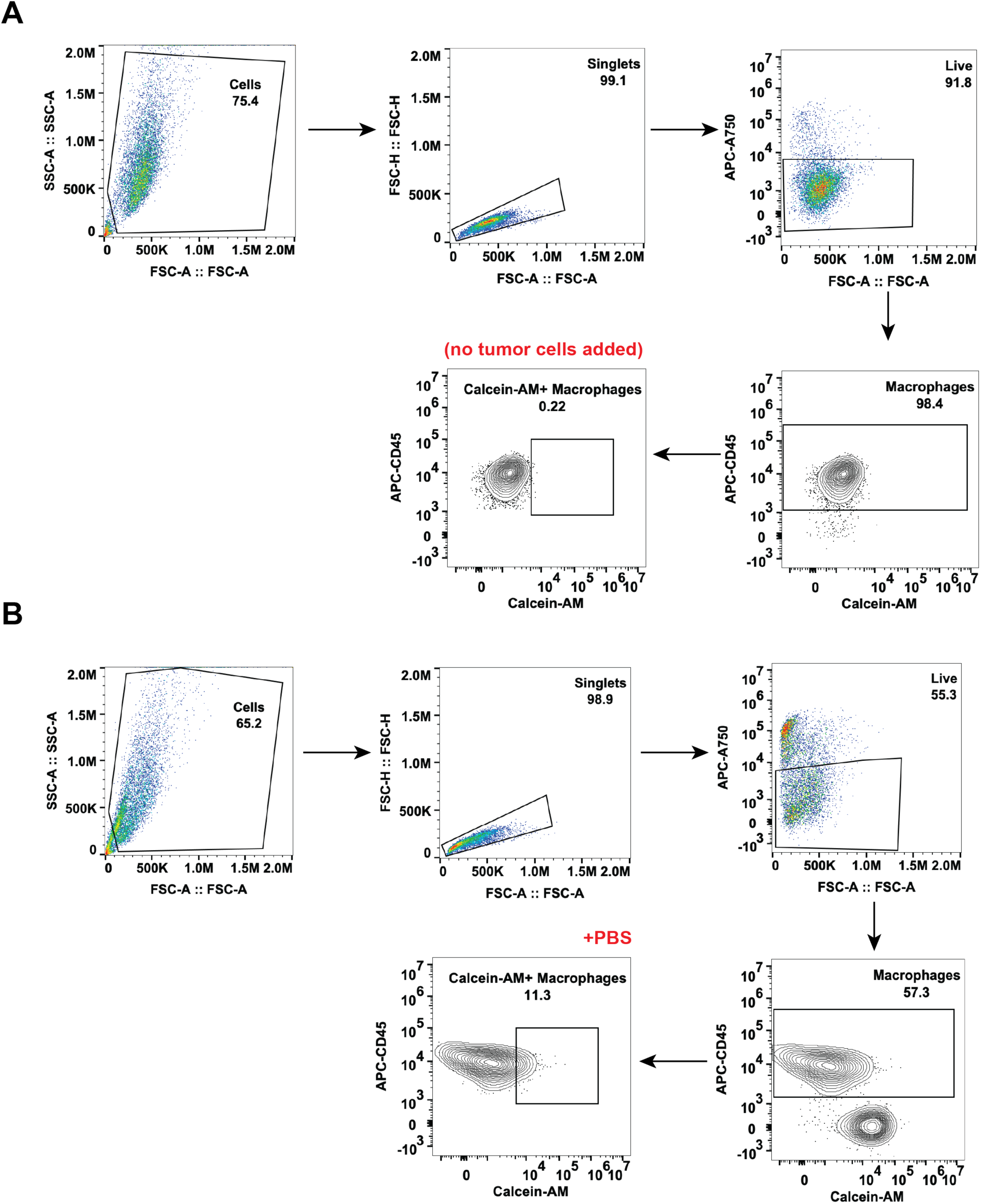
Representative flow-cytometry gating tree used in phagocytosis assays. (A) To identify macrophages, singlets were gated using FSC-A/FSC-H and dead cells were excluded using Zombie NIR fixable viability kit. CD45^+^ cells were considered macrophages. (B) Macrophages were incubated with calcein-AM stained DMG cells. Single-stained samples were used to identify the correct gate for macrophages (CD45^+^) and tumor cells (Calcein-AM^+^). The CD45^+^ and Calcein-AM^+^ population represents macrophages that have successfully phagocytosed DMG cells.

**Supplementary Figure 4.**
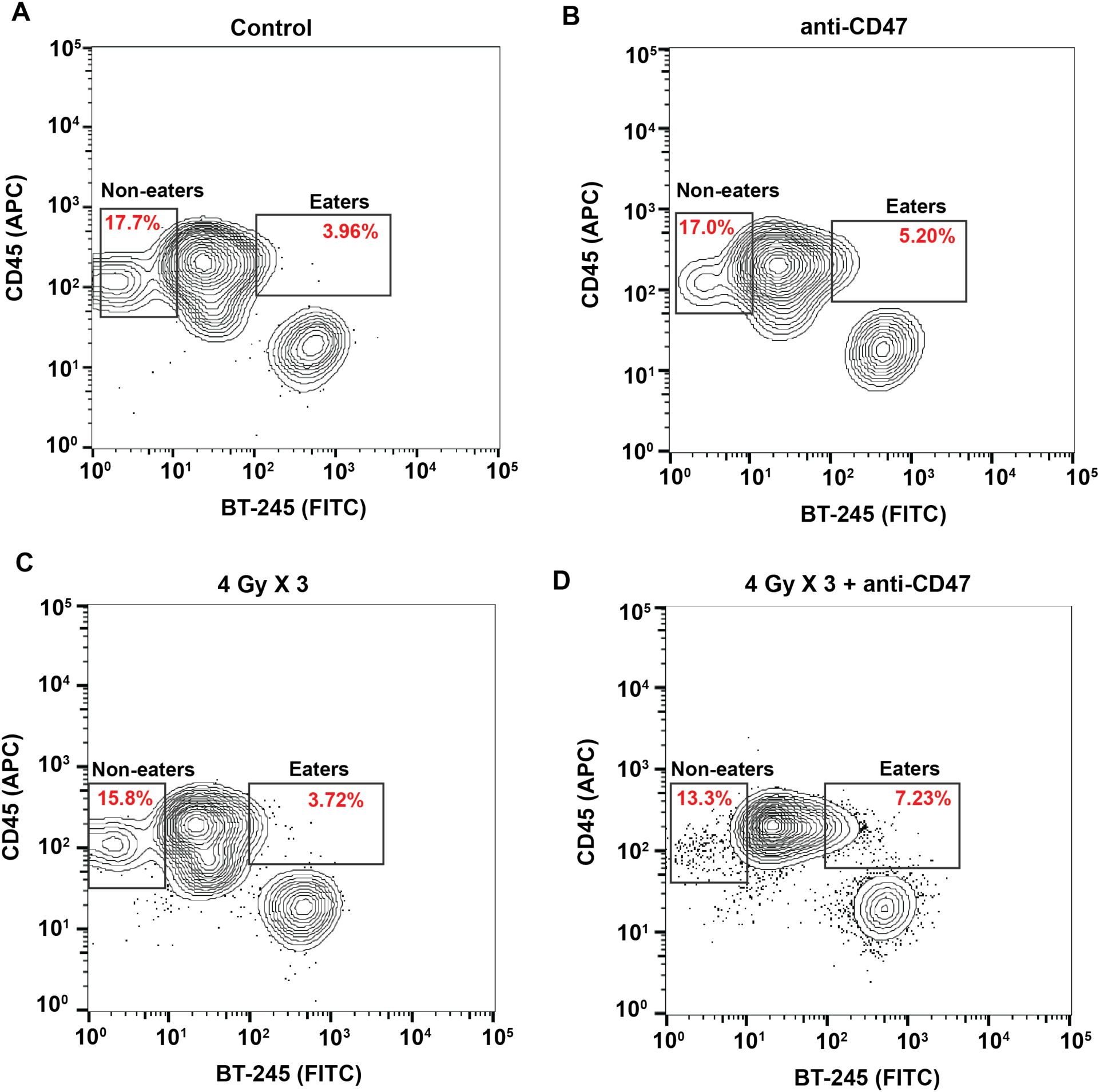
Flow cytometry gating strategy to isolate “eaters” and “non-eaters” for single-cell RNA-sequencing. (A-D) Macrophages incubated with B245 cells pretreated with either anti-CD47 therapy, radiation therapy (RT: 4 Gy X 3), or a combination (4 Gy X 3 + anti-CD47) of both and stained with calcein-AM. Macrophages incubated with tumor cells that were not pretreated was used as control. Macrophages that phagocytosed (“eaters;” CD45^+^ and Calcein-AM^+^) or those did not (“non-eaters; CD45^+^ only) tumor cells were identified and isolated using flow cytometry.

**Supplementary Figure 5.**
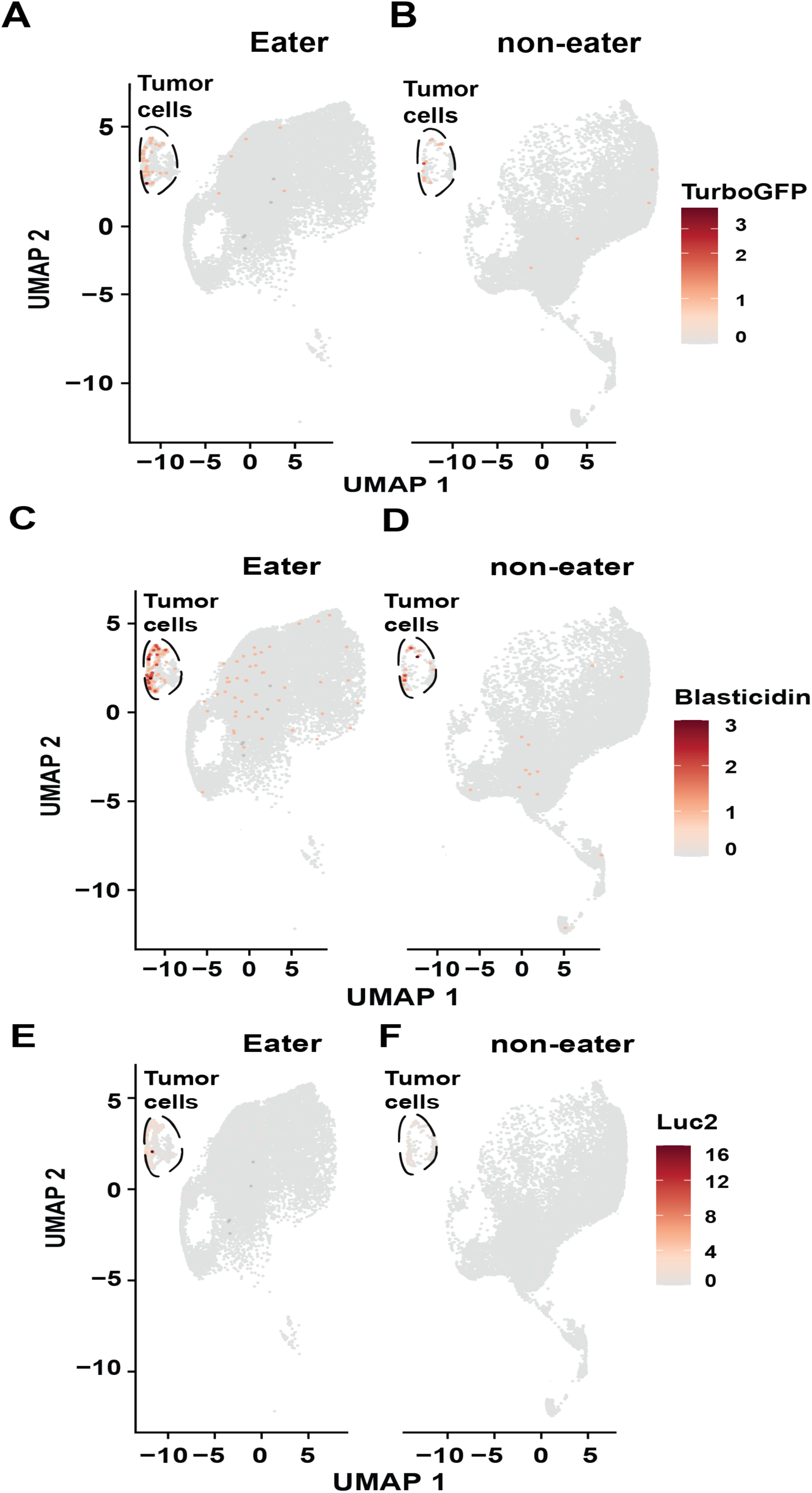
Identification of tumor cells in the single-cell RNA-seq dataset. UMAP dimensionality reduction plots show the distribution of tumor cells and macrophages. Annotation of the tumor cells (indicated by dotted lines) using the specific marker of green fluorescent protein (GFP) **(A & B)**, blasticidin **(C & D)** and firefly luciferase reporter **(E & F)**.

**Supplementary Figure 6.**
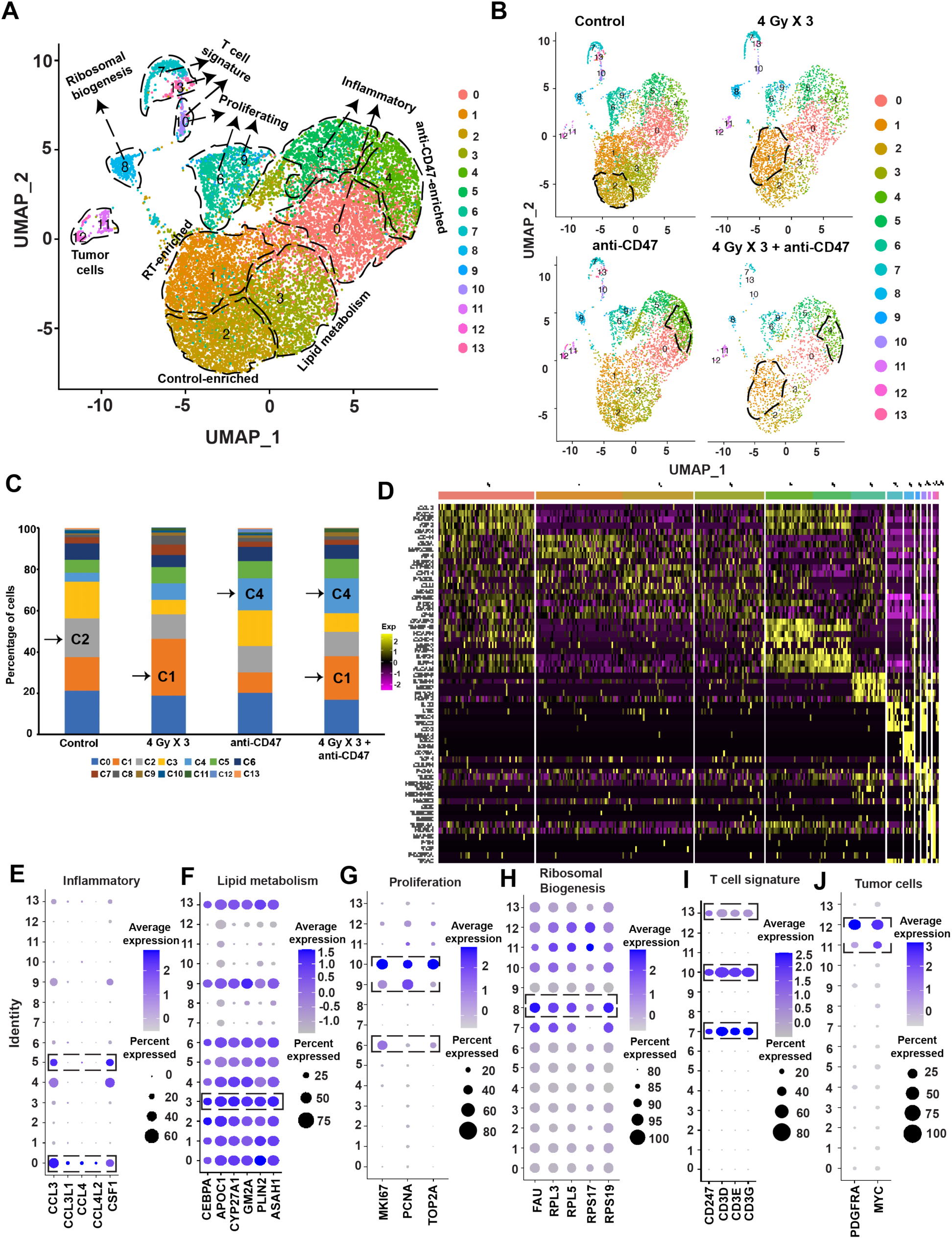
Single-cell RNA-sequencing reveals the enrichment of distinct macrophage subsets following co-culture with either Control, RT, anti-CD47 or RT+ anti-CD47 treated DMG cells. (A) UMAP projection displaying 13 distinct cell clusters from the non-eater’s cohort. Each dotted line and arrow indicate the identity of that cell cluster. **(Band C)** UMAP plots and proportion of cells in each cluster from 4 treatment groups: control, 4 Gy X 3, anti-CD47 or 4 Gy X 3 + anti-CD47. Dotted lines indicate the expansion/enrichment of distinct macrophage cluster in that treatment condition. Note the expansion/enrichment of two distinct cell clusters (1 and 4) in the combination treatment. **(D)** Heatmap of marker genes in each cell cluster. Representative genes are highlighted on the left. Bubble plots demonstrating expression of marker genes associated with inflammatory **(E)**, lipid metabolism **(F)**, Proliferation **(G)**, Ribosomal Biogenesis **(H)**, T cell signature **(I)**, and tumor cell signature **(J)** by the various cell clusters. The dotted box indicates the cell clusters with higher average marker gene expression. The size of the bubble dot is proportional to the percentage of cells in a cluster expressing the marker gene and the color intensity is proportional to average scaled marker gene expression within a cluster.

**Supplementary Figure 7.**
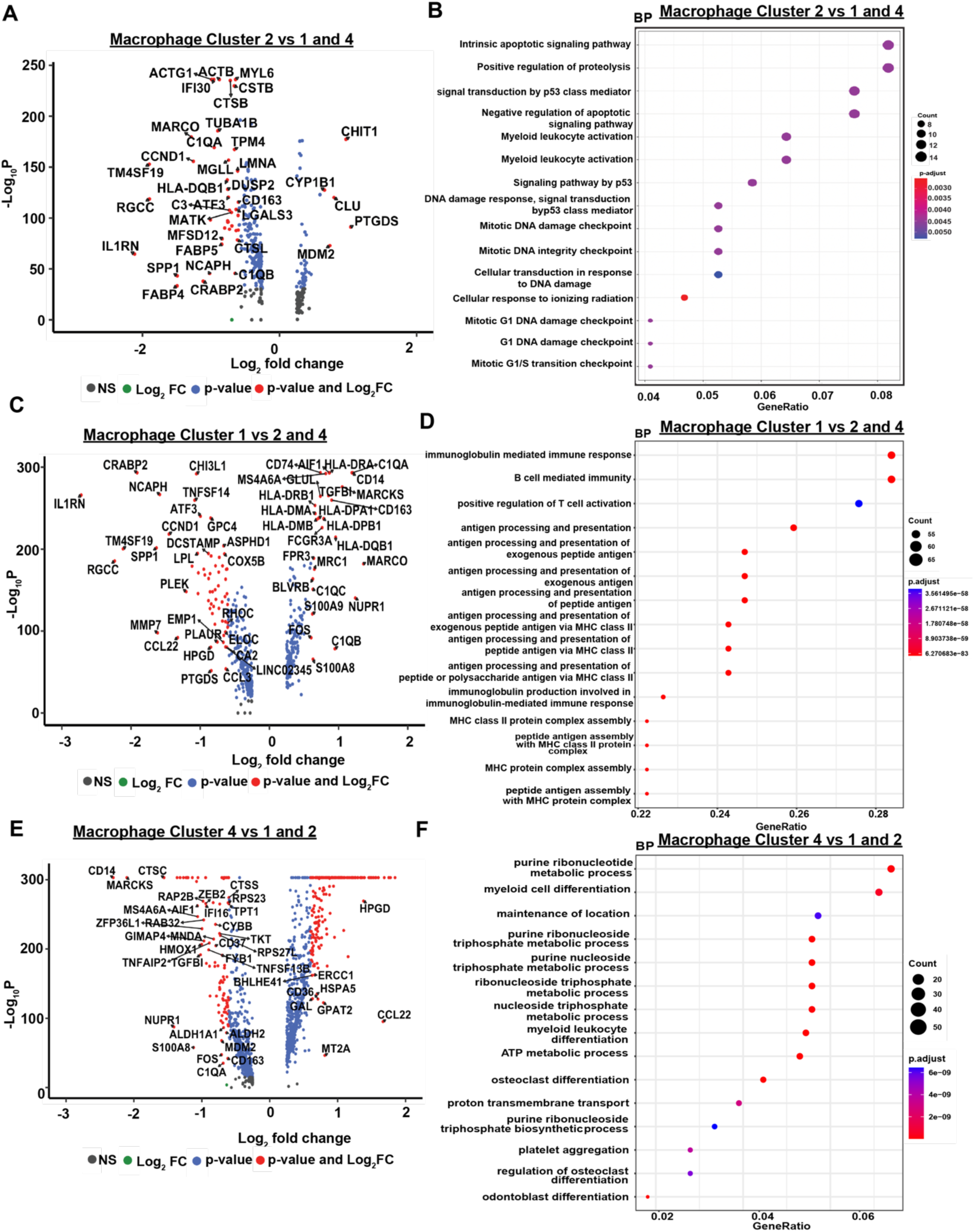
Characterization of macrophages that didn’t phagocytose diffuse midline glioma cells pretreated with Control, anti-CD47 therapy or RT. **(A)** Volcano plot showing the top differentially upregulated and downregulated genes in the macrophages that are enriched after co-culture with control treated BT245 cells (cell cluster 2) compared to macrophages that were enriched after co-culture with either anti-CD47 (cell cluster 1) or RT (cell cluster 4) treated BT245 cells. (**B**) Gene Ontology enrichment analysis of biological process for significantly upregulated genes between control-enriched macrophages vs anti-CD47- and RT- enriched macrophages. Note only the top 15 biological processes are shown. (**C)** Volcano plot showing the top differentially upregulated and downregulated genes in the macrophages that are enriched after co-culture with anti-CD47 treated BT245 cells (cell cluster 1) compared to macrophages that were enriched after co-culture with either control (cell cluster 2) or RT (cell cluster 4) treated BT245 cells. (**D**) Gene Ontology enrichment analysis of biological process for significantly upregulated genes between anti-CD47-enriched macrophages vs Control- and RT- enriched macrophages. Note only the top 15 biological processes are shown. **(E)** Volcano plot showing the top differentially upregulated and downregulated genes in the macrophages that are enriched after co-culture with RT treated BT245 cells (cell cluster 4) compared to macrophages that were enriched after co-culture with either control (cell cluster 1) or anti-CD47 (cell cluster 2) treated BT245 cells. (**F)** Gene Ontology enrichment analysis of biological process for significantly upregulated genes between control-enriched macrophages vs anti-CD47- and RT- enriched macrophages. Note only the top 15 biological processes are shown.

**Supplementary Figure 8.**
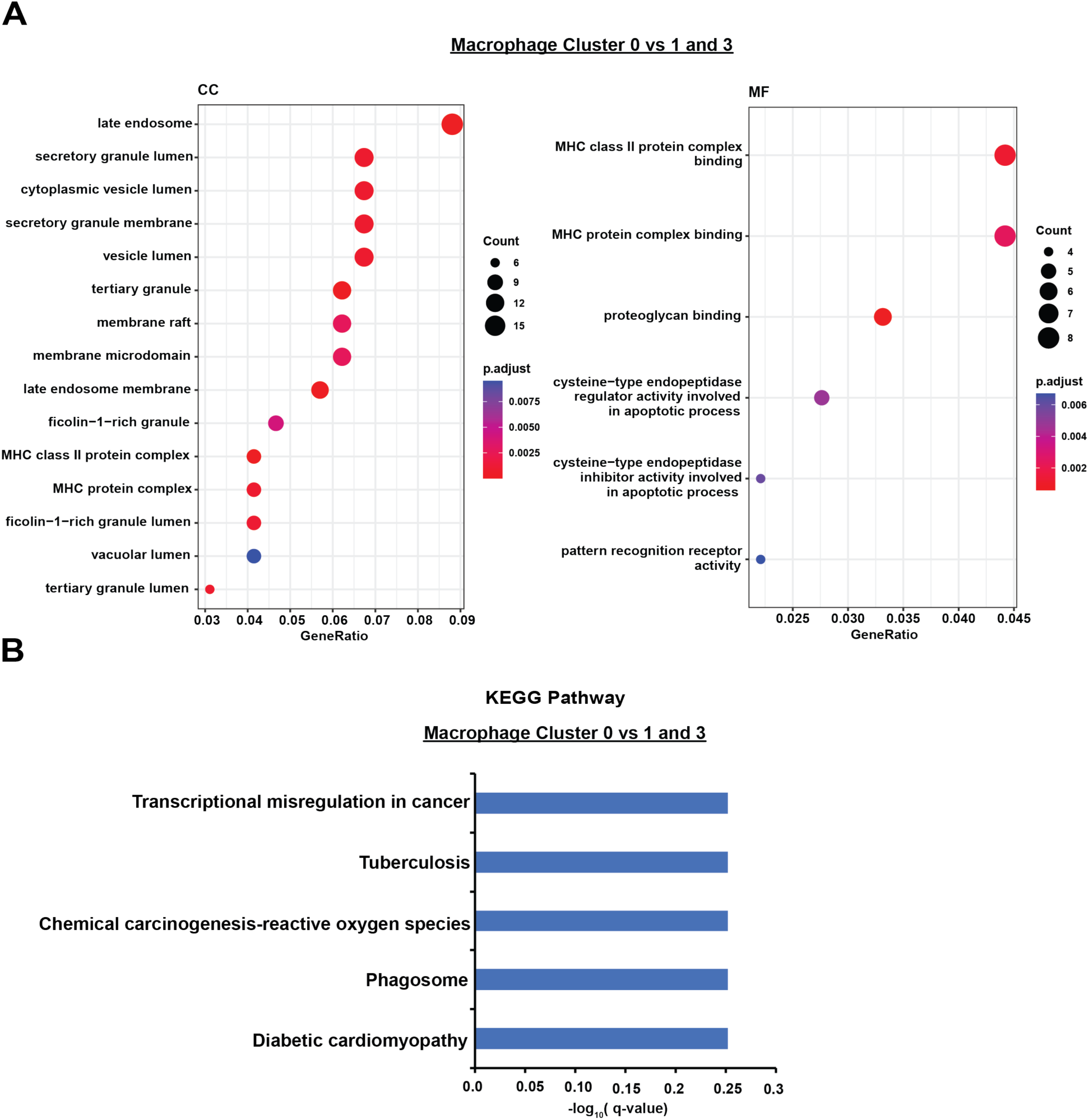
Gene set enrichment and KEGG pathway analysis of differentially expressed genes identified in macrophages enriched under control condition compared to those enriched under anti-CD47 or RT treated condition in the “eater’s” dataset.

**Supplementary Figure 9.**
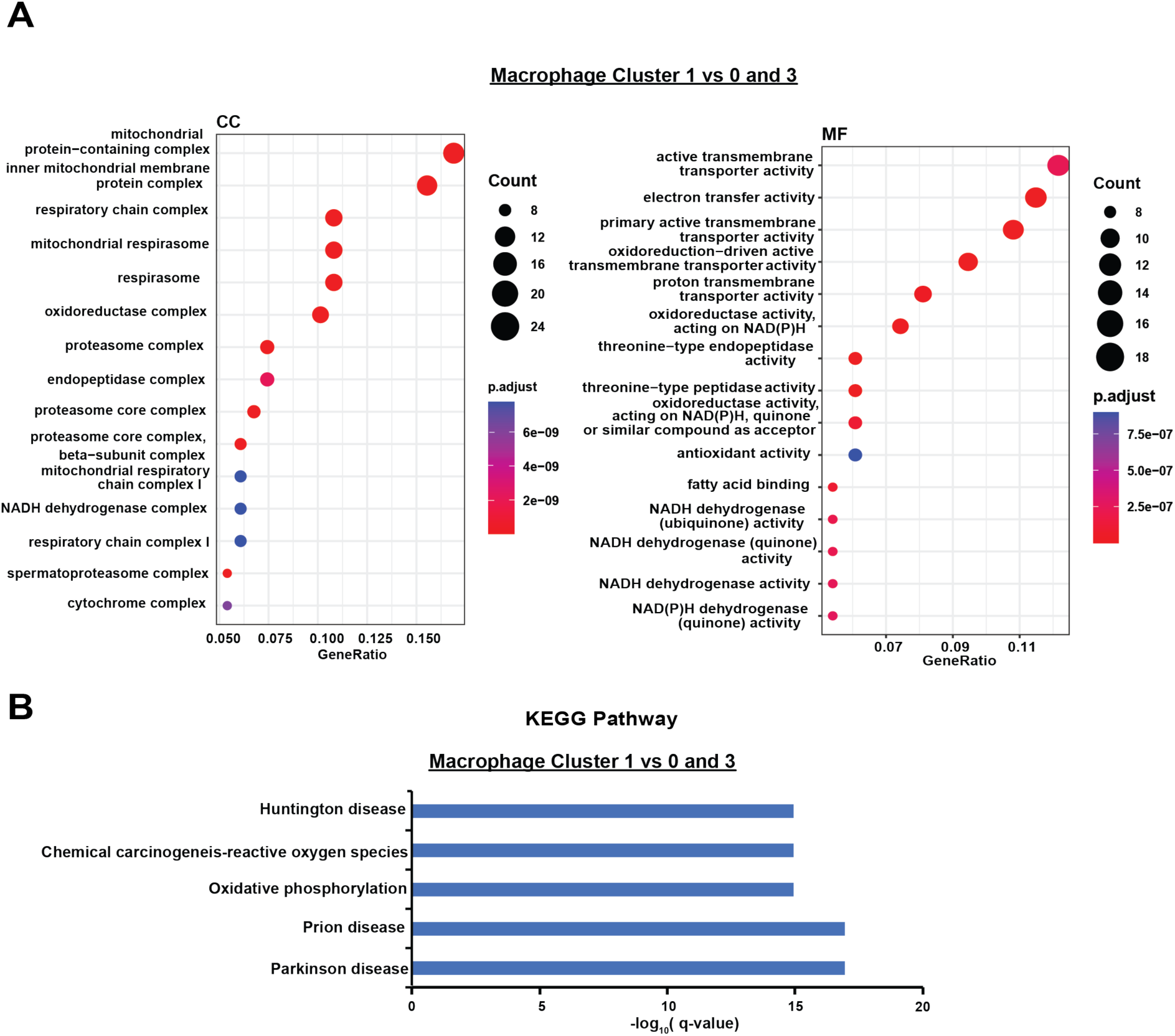
Gene set enrichment and KEGG pathway analysis of differentially expressed genes identified in macrophages enriched under anti-CD47 condition compared to those enriched under control or RT treated condition in the “eater’s” dataset.

**Supplementary Figure 10.**
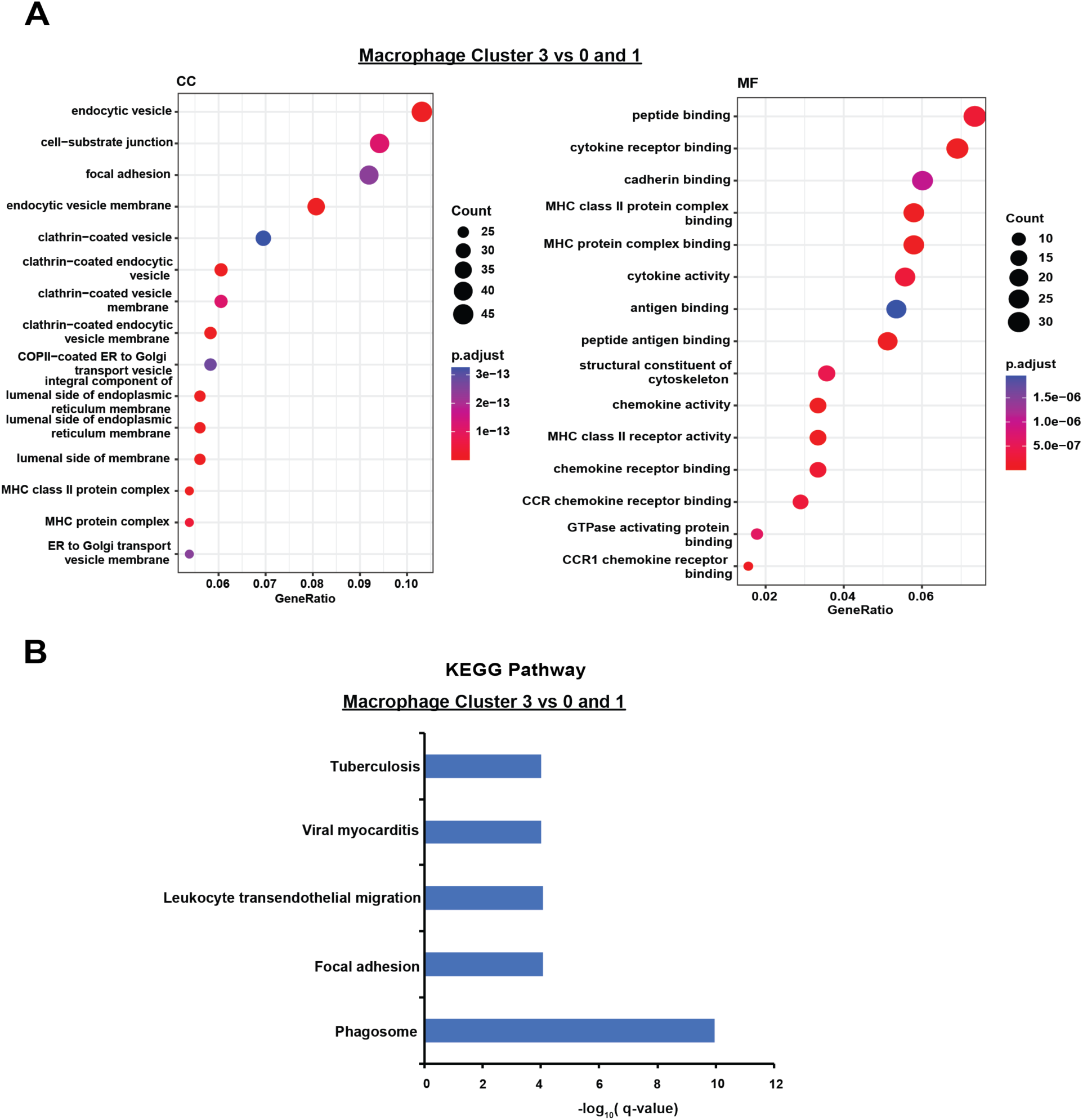
Gene set enrichment and KEGG pathway analysis of differentially expressed genes identified in macrophages enriched under RT condition compared to those enriched under control or anti-CD47 treated condition in the “eater’s” dataset.

**Supplementary Figure 11.**
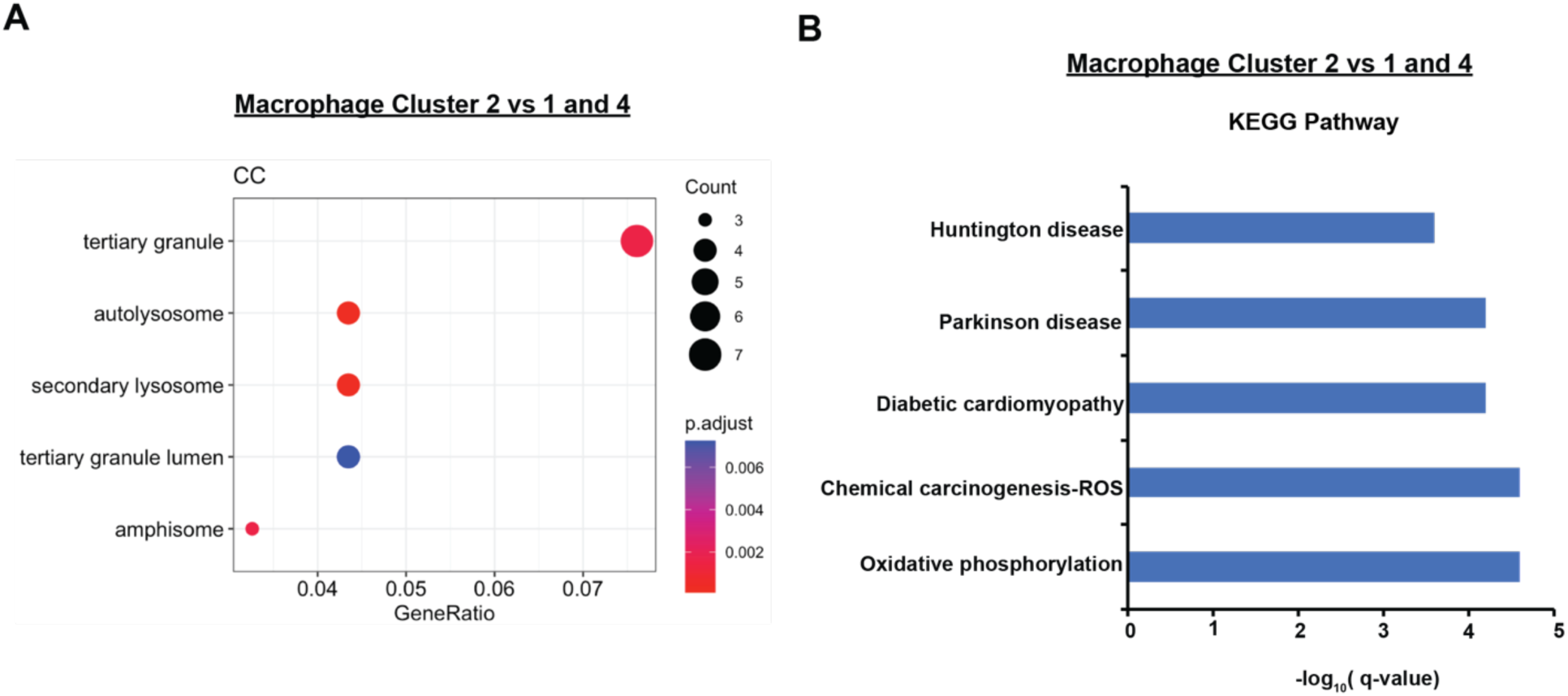
Gene set enrichment and KEGG pathway analysis of differentially expressed genes identified in macrophages enriched under control condition compared to those enriched under RT or anti-CD47 treated condition in the “non-eater’s” dataset.

**Supplementary Figure 12.**
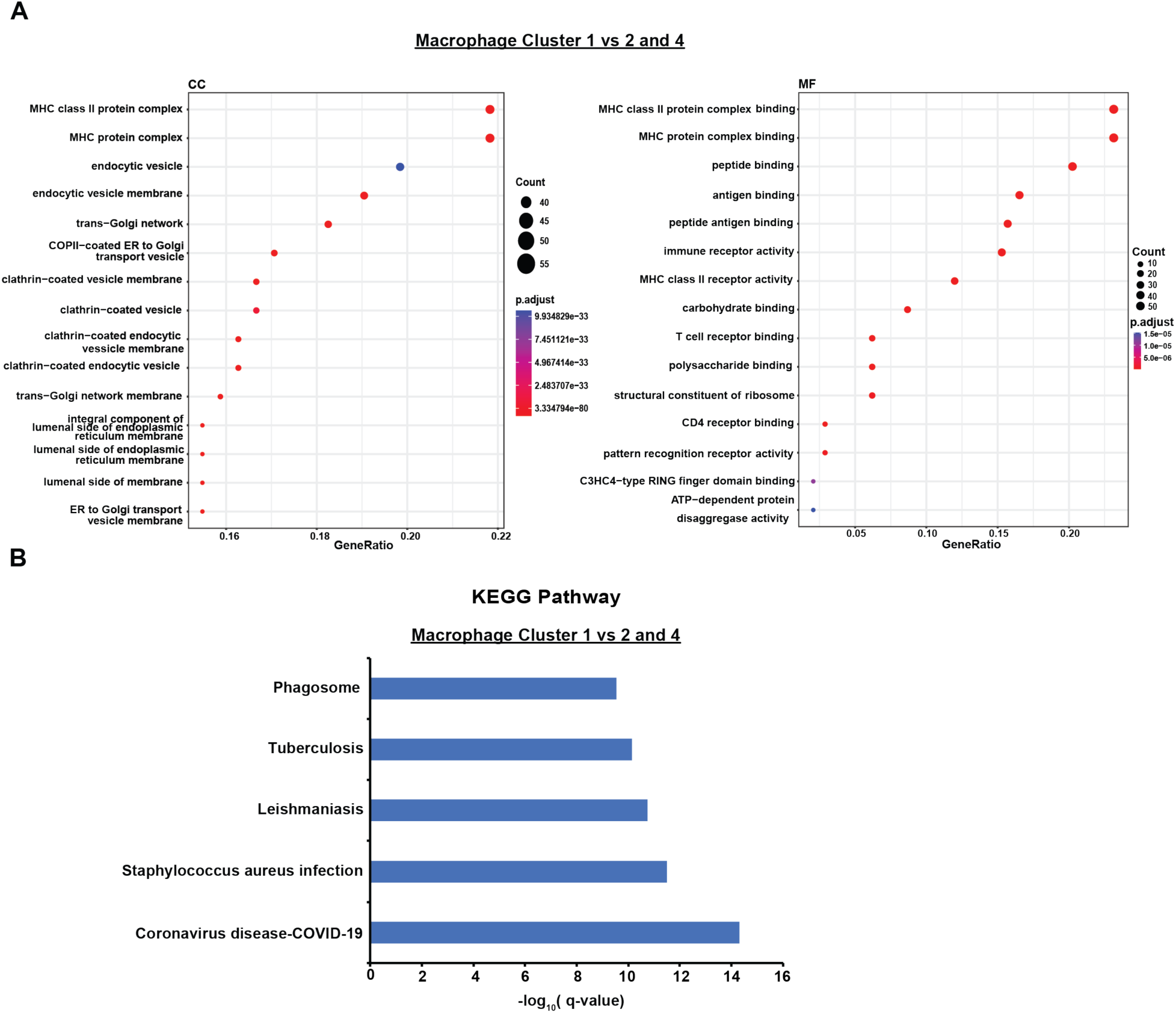
Gene set enrichment and KEGG pathway analysis of differentially expressed genes identified in macrophages enriched under RT condition compared to those enriched under control or anti-CD47 treated condition in the “non-eater’s” dataset.

**Supplementary Figure 13.**
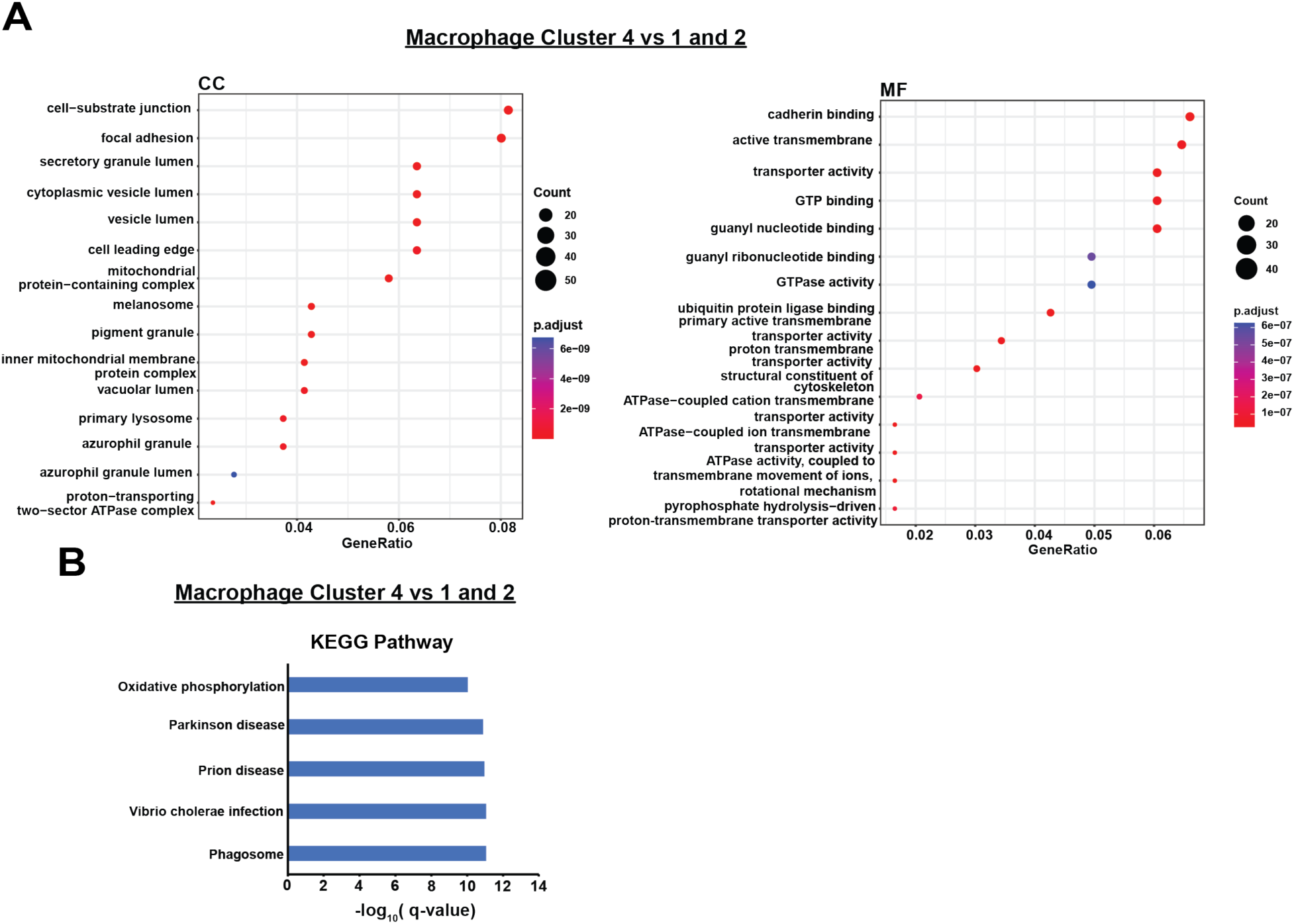
Gene set enrichment and KEGG pathway analysis of differentially expressed genes identified in macrophages enriched under anti-CD47 condition compared to those enriched under control or RT treated condition in the “non-eater’s” dataset.

**Supplementary Figure 14.**
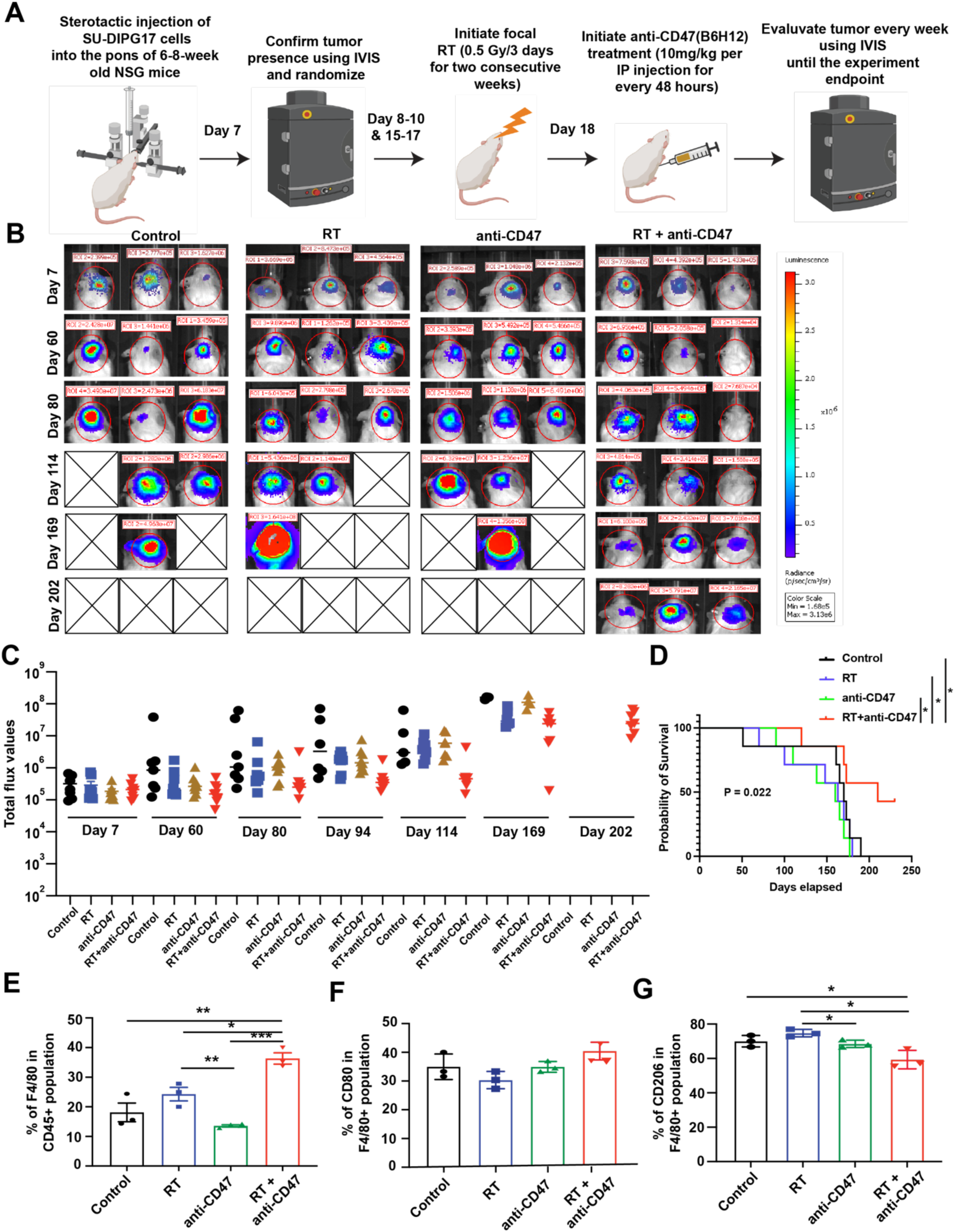
Combination of RT and anti-CD47 treatment reduces tumor burden and prolongs the survival of mice-bearing DIPG17 xenografts compared to monotherapy. **(A)** Schematic diagram showing the experimental treatment plan followed**. (B)** Representative bioluminescence images of the mice-bearing Luc2-expressing DIPG17 cells before, after, and during the specified treatments. The scale bar adjacent to the image displays bioluminescence counts (photons/second/cm^2^/steradian). **(C)** Quantification of total IVIS flux values over time course. **(D)** Kaplan-Meier survival analysis of BT245 xenografts with indicated treatments, control, n = 7; RT, n= 7; anti-CD47, n = 7; and RT + anti-CD47, n = 9. The log-rank test was used to calculate statistical significance. *p < 0.05, **p < 0.01. **(E-G)** Bar graphs demonstrating the relative percentages of F4/80^+^, CD80^+^ (M1-like) and CD206^+^ (M2-like) tumor-associated macrophages in control, RT, anti-CD47, or RT + anti-CD47 treated mice-bearing DIPG17 xenografts. Data shown are obtained from n=3 mice for each group and are represented as mean ± SD. Unpaired student’s t-test: *p < 0.05, **p < 0.01 and ***p < 0.0001.

**Supplementary Figure 15.**
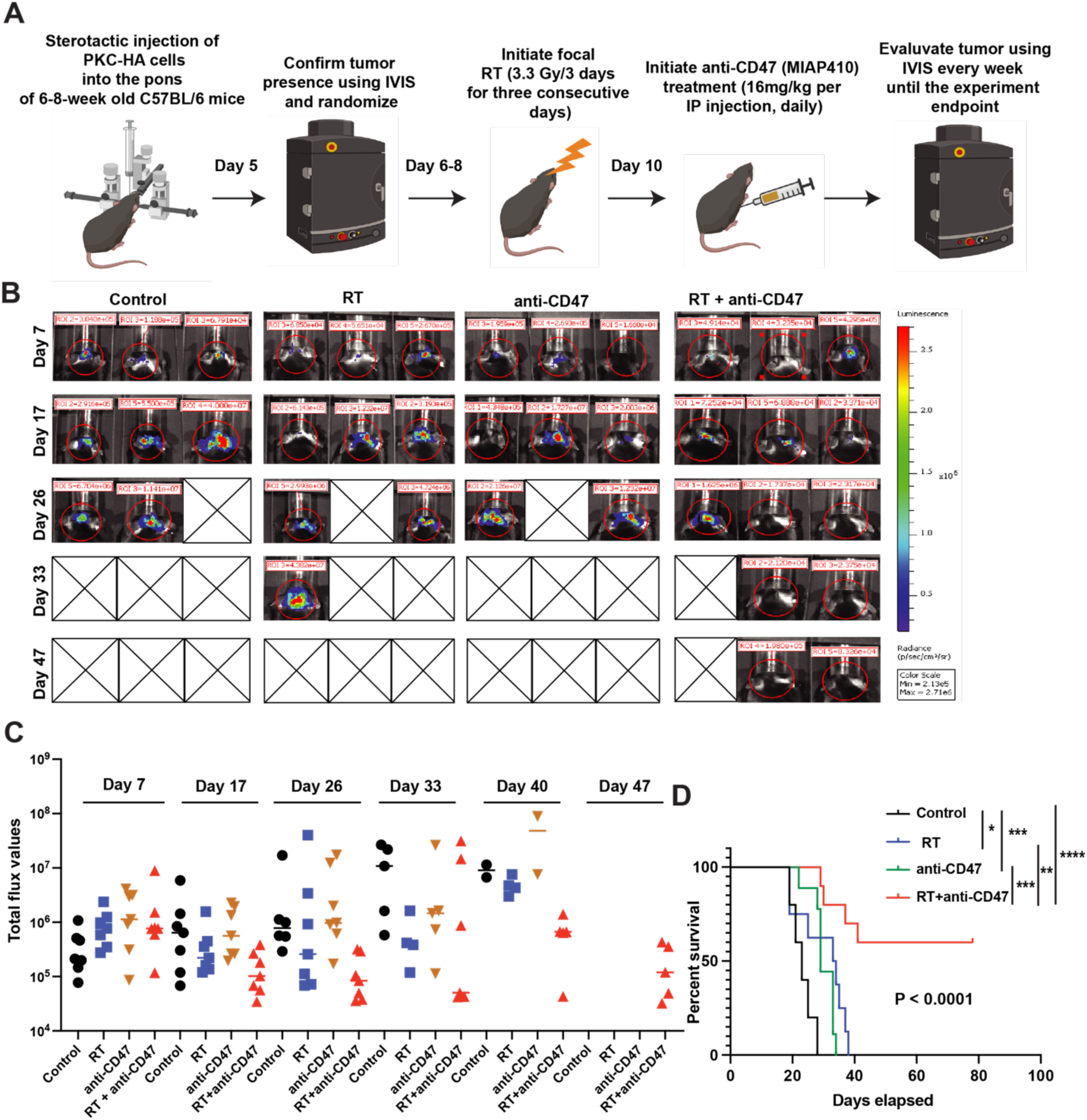
Combination of RT and anti-CD47 treatment reduces tumor burden and prolongs the survival of mice-bearing PKC-HA xenografts compared to monotherapy. **(A)** Schematic diagram showing the experimental treatment plan followed**. (B)** Representative bioluminescence images of the mice-bearing Luc2-expressing PKC-HA cells before, after, and during the specified treatments. The scale bar adjacent to the image displays bioluminescence counts (photons/second/cm^2^/steradian). **(C)** Quantification of total IVIS flux values over time course. **(D)** Kaplan-Meier survival analysis of BT245 xenografts with indicated treatments, control, n = 7; RT, n = 7; anti-CD47, n = 7; and RT + anti-CD47, n=7. The log-rank test was used to calculate statistical significance. *p < 0.05, **p < 0.01. Data shown is obtained from n=3 mice for each group and are represented as mean ± SD.

**Supplementary Figure 16.**
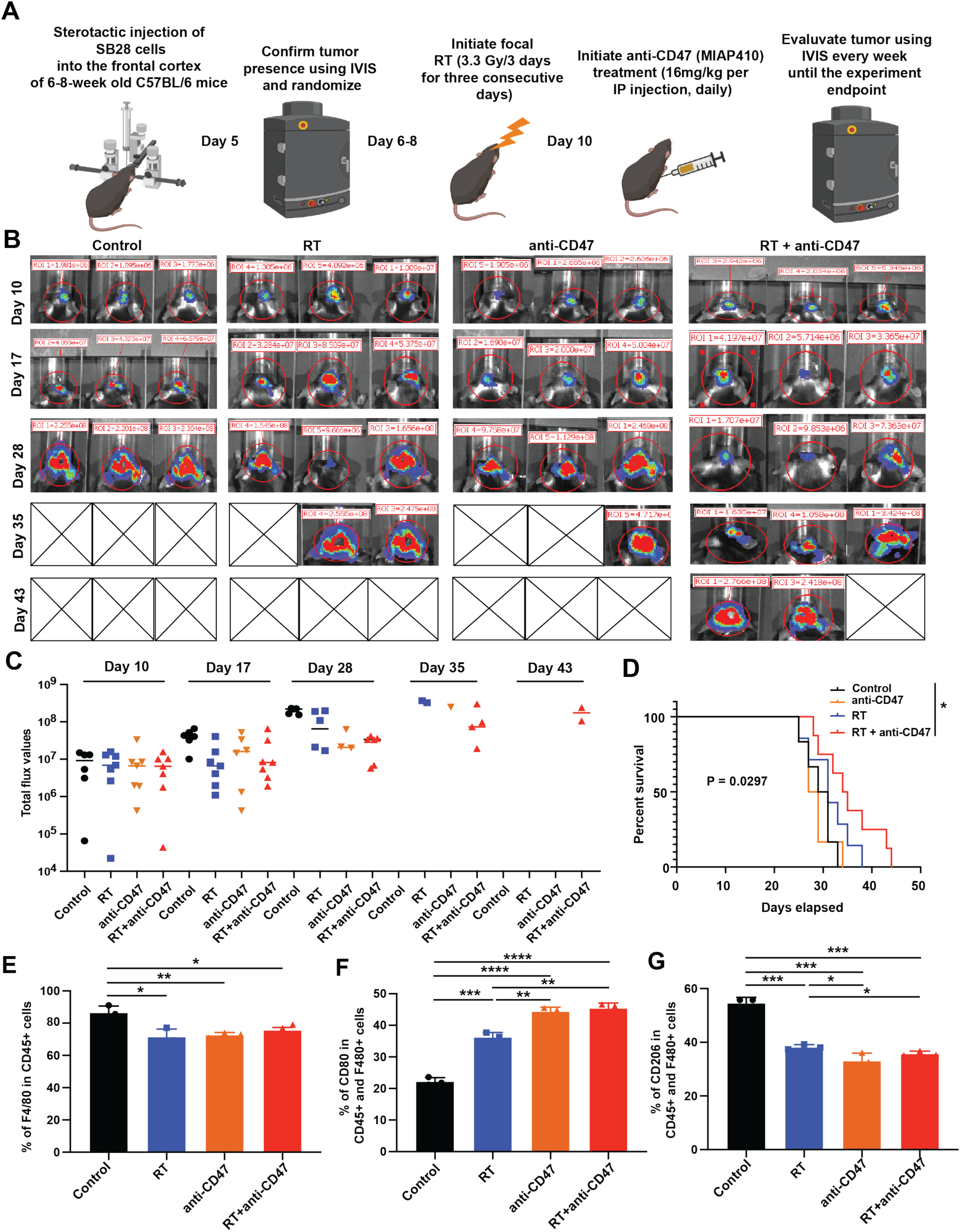
Combination of RT and anti-CD47 treatment reduces tumor burden and prolongs the survival of mice-bearing SB28 xenografts compared to monotherapy. **(A)** Schematic diagram showing the experimental treatment plan followed**. (B)** Representative bioluminescence images of the mice-bearing Luc2-expressing SB28 cells before, after, and during the specified treatments. The scale bar adjacent to the image displays bioluminescence counts (photons/second/cm^2^/steradian). **(C)** Quantification of total IVIS flux values over time course. **(D)** Kaplan-Meier survival analysis of BT245 xenografts with indicated treatments, control, n = 10; RT, n= 9; anti-CD47, n=10; and RT + anti-CD47, n=10. The log-rank test was used to calculate statistical significance. *p < 0.05, **p < 0.01. **(E-G)** Bar graphs demonstrating the relative percentages of F4/80^+^, CD80^+^ (M1-like) and CD206^+^ (M2-like) tumor-associated macrophages in control, RT, anti-CD47, or RT + anti-CD47 treated mice-bearing SB28 xenografts. Data shown are obtained from n=3 mice for each group and are represented as mean ± SD. Unpaired student’s t-test: *p < 0.05, **p < 0.01 and ***p < 0.0001.

